# Impacts of Simultaneous Multislice Acquisition on Sensitivity and Specificity in fMRI

**DOI:** 10.1101/243782

**Authors:** Benjamin B. Risk, Mary C. Kociuba, Daniel B. Rowe

## Abstract

Simultaneous multislice (SMS) acquisition can be used to decrease the time between acquisition of fMRI volumes, which can increase sensitivity by facilitating the removal of higher-frequency artifacts and boosting effective sample size. The technique requires an additional processing step in which the slices are separated, or unaliased, to recover the whole brain volume. However, this may result in signal “leakage” between aliased locations, i.e., slice “leakage,” and lead to spurious activation (decreased specificity). SMS can also lead to noise amplification, which can reduce the benefits of decreased repetition time. In this study, we evaluate the original slice-GRAPPA (no leak block) reconstruction algorithm and acquisition scenarios used in the young adult Human Connectome Project (HCP), as well as split slice-GRAPPA (leak block). In addition to slice leakage, signal leakage can result from spatial smoothing, i.e., smoothing leakage, which leads to inflated regions of activation. Previous studies have generally found that SMS acquisition results in higher test statistics and/or a greater number of activated voxels. Here, we use simulations to disentangle this phenomenon into true positives (sensitivity) and false positives (decreased specificity). Slice leakage was greatly decreased by split slice-GRAPPA. Noise amplification was decreased by using moderate acceleration factors (AF = 4). We examined slice leakage in unprocessed fMRI motor task data from the HCP, which used the original slice-GRAPPA. When data were smoothed, we found evidence of slice leakage in some, but not all, subjects. We also found evidence of SMS noise amplification in unprocessed task and processed resting-state HCP data.

## 1 Introduction

The rapid evolution in simultaneous multislice (SMS) imaging techniques has led to large decreases in image acquisition time, which has increased the temporal resolution in functional magnetic resonance imaging (fMRI) and improved tractography in diffusion MRI. In SMS imaging, a multiband radio frequency (RF) pulse with slice-selective gradient is utilized to simultaneously collect data from multiple slices (Feinberg and Setsompop, 2013). The time to repetition (TR) is decreased in proportion to the number of simultaneously excited slices, and a twelve-fold reduction has been implemented for 2D echo-planar imaging (EPI) (Xu et al., 2013). Higher sampling rates can improve the ability to separate physiological artifacts such as breathing and cardiac pulsation, which contain power at higher frequencies than the hemodynamic response function (Smith et al., 2013; Boyacioğlu et al., 2015; Griffanti et al., 2014). Test statistics and temporal signal-to-noise ratio (tSNR) may benefit from an increase in the effective sample size resulting from the collection of a higher number of volumes in the same run duration (Todd et al., 2016; Feinberg et al., 2010). SMS techniques are becoming increasingly important because they are crucial to decreasing acquisition times in 7T MRI, which decreases geometric distortion from inhomogeneity effects at ultrahigh magnetic fields, and thus serve a central role in high-resolution brain mapping (Moeller et al., 2010; Poser and Setsompop, 2017; Uğurbil et al., 2013).

The benefits of SMS are in part offset by noise amplification due to slice separation during reconstruction. Additionally, signal loss and poor contrast can occur with high acceleration factors (AFs) when the TR is less than the T1 relaxation time of the tissue of interest (Smith et al., 2013). The specific absorption rate is also a concern with SMS, as power deposition from the multiband (MB) RF pulses increases proportionally to the number of slices simultaneously excited. Noise amplification can be quantified using coil geometry factors, or *g*-factors. *G*-factors can be improved by exploiting spatial variation in coil sensitivities. In single-shot 2D EPI, gradient blips are applied to slices collected at the same time to achieve field of view shifts to improve *g*-factors (Breuer et al., 2005; Nunes et al., 2006; Setsompop et al., 2012). SMS protocols using FOV/3 shifts and an AF equal to eight have been popularized by the Human Connectome Project (HCP) acquisition protocol (Glasser et al., 2016), and SMS imaging is being used in other major studies such as the UK Biobank (Miller et al., 2016) and Rhineland Study (Breteler et al., 2014). Advanced methods are being developed for 3D encoding that more efficiently utilize 3D variation in coil sensitivities using corkscrew-like trajectories (Bilgic et al., 2015).

In addition to noise amplification, the accuracy of image reconstruction in SMS is impacted by slice leakage, or signal from aliased slices that is not adequately separated from a given slice (Moeller et al., 2012). Slice leakage can be difficult to detect in SMS imaging, where differences in the accuracy of kernels can result in one slice showing significant aliasing while another slice is free from artifact (Barth et al., 2016). The *L*-factor can be quantified from single-band training data, where it is equal to the image reconstructed when applying the GRAPPA kernel from a focal slice to the aliased slices (excluding the focal slice) (Cauley et al., 2014). *L*-factors generally increase with AF and are oftentimes lower in blipped-CAIPI acquisition protocols versus non-blipped (Xu et al., 2013).

Previous empirical studies have interpreted higher test statistics and/or a larger number of activated voxels in SMS data versus single-band data as evidence of increased sensitivity, although this is not equivalent to statistical sensitivity because the truth is unknown. Higher test statistics in the presence of bias, for example due to slice leakage or spatial smoothing, result in a decrease in specificity. Higher test statistics and/or larger spatial extents of activation is SMS data have been found in task fMRI (Chen et al., 2015; Boyacioğlu et al., 2015; Demetriou et al., 2016; Todd et al., 2016, 2017) and resting-state fMRI (Preibisch et al., 2015; Feinberg et al., 2010; Smith et al., 2013). Approaches comparing test statistics can also be hindered by inaccurate modeling of the time-series errors for shorter TRs, and the use of low-order AR(1) models may lead to inflated test statistics (Chen and Glover, 2017) (but see Todd et al. (2017), who account for this issue by down sampling).

While previous studies address the potential benefits of SMS, the costs in terms of false positives have been less studied. Todd et al. (2016) developed an empirical formulation of specificity by examining activation in voxels aliased to voxels believed to reflect true activation in visual and motor tasks. For a given AF, a voxel was deemed a false positive if it was activated and aliased to the voxel with largest test statistic from an activation cluster but was inactive in the other runs collected using a different AF (in which the aliased locations differed). Their study compared the incidence of suspected false positives in reconstruction using the original slice-GRAPPA (no leak block) (Setsompop et al., 2012) and split slice-GRAPPA (leak block) (Cauley et al., 2014). Split slice-GRAPPA dramatically reduced the instances of false positives without reducing the size of the top 1% of test statistics. The authors’ approach focused on single voxels with the largest test statistics, whereas researchers may also be interested in whether slice leakage can lead to clusters of voxels that can be mistaken to be of biological origin. Additionally, the study evaluated through-plane multiband factors in combination with 2× in-plane acceleration. The HCP young adult study and UK Biobank use AF = 8 without in-plane acceleration, and the incidence of false positives has not been evaluated for their acquisition protocols. Finally, the study used a small amount of smoothing (2 mm FWHM), and the interaction between AF and smoothing has not been examined.

The questions and insights raised by Todd et al. (2016) in part motivated our investigation of the patterns of *regional* aliasing and a formal evaluation of the statistical impacts of SMS acquisition including the impacts of smoothing. The impacts of signal leakage have been demonstrated for in-plane acceleration, where interpolation using the in-plane GRAPPA kernel and spatial smoothing induce correlations between aliased and neighboring voxels (Bruce and Rowe, 2014). Here, we characterize signal leakage due to *slice* leakage, which causes signal to leak into distant voxels, and *smoothing* leakage, which is associated with signal leaking into neighboring voxels. We define regional aliasing as a set of contiguous voxels aliased to a region of interest. Slice leakage can result in a cluster of activation in regionally aliased voxels in task fMRI, or in a cluster of induced correlations in resting-state fMRI. The costs and benefits of SMS can be quantified by statistical sensitivity and specificity. Statistical sensitivity is the true positive rate, and in the context of hypothesis testing, it is equal to the proportion of rejected null hypotheses given the null hypothesis is false (this is also called power, and is equivalent to one minus the probability of a false negative, i.e., one minus the type II error rate). Specificity is the true negative rate, which is the proportion of null hypotheses that were not rejected given the null hypothesis is true (this is is equal to one minus the probability of a false positive, i.e., one minus the type I error rate). Disentangling the impacts of higher test statistics into true positives and false positives requires simulations where the truth is known.

The goal of this study is to characterize the trade offs in SMS in terms of sensitivity and specificity, with a focus on task fMRI. Large *g*-factors, low tSNR, and large *L*-factors are detrimental to image reconstruction, but it is difficult to determine tolerable levels of these measures and translate them into statistical outcomes for experimental design. Whereas studies in MRI acquisition often focus on *g*-factors, here we examine the standard deviation of the residuals from task-activation models and the standard deviation of time series from resting-state data, which are closely related to *g*-factors but may be more familiar to behavioral scientists. Rather than focus on *L*-factors, we examine patterns of regional aliasing, which has practical implications for activation studies. We evaluate how high multiband factors can influence sensitivity and specificity for a left-hand motor task experiment motivated by the acquisition protocols, reconstruction algorithm, and task data from the Human Connectome Project. One of the primary aims of the HCP is the development of MRI acquisition protocols with tested and optimized pulse sequences and image reconstruction algorithms (Glasser et al., 2016). In this work, we evaluate the original slice-GRAPPA (no leak block) algorithm from Setsompop et al. (2012), which was used in the HCP young adult study and UK Biobank. We also evaluate split slice-GRAPPA (leak block) in simulations. Our goal is to provide information that can aid experimenters implementing the HCP’s acquisition protocol. A second goal is to provide information to scientists analyzing and interpreting HCP and UK Biobank data.

In the next section, we propose a technique that allows researchers to examine whether spurious activation is occurring in brain regions aliased with areas where prior knowledge indicates true activation. In Section 3, we conduct a simulation study. In Section 4, we analyze HCP motor task data for evidence of slice leakage and SMS noise amplification. In Section 5, we examine noise amplification in the ICA-FIX preprocessed resting-state data.

## 2 Regional aliasing in Slice-GRAPPA

Before describing how aliasing patterns can be calculated, we provide background on controlled aliasing in SMS. The *g*-factor was introduced for in-plane parallel imaging with sensitivity-based (SENSE) reconstruction (Pruessmann et al., 1999). The *g*-factor in parallel imaging can be defined as the ratio of the signal-to-noise ratio (SNR) for the unaccelerated data to the SNR of the accelerated data divided by the square-root of the under-sampling rate, or assuming the signals are equivalent, the ratio of the standard deviations of the accelerated data to the unaccelerated data divided by the square-root of the under-sampling rate (Robson et al., 2008). In the acquisition scenarios considered in our study, there is no in-plane acceleration and a measure of the *g*-factor is the ratio of the standard deviations of the SMS to single-band data. To decrease *g*-factors, CAIPIRINHA techniques apply phase modulations to achieve shifts in the field of view, which allow the variation in coil strengths to be exploited for improved slice separation. CAIPIRINHA was developed for multi-shot sequences (Breuer et al., 2005), but was not applicable to EPI acquisitions (diffusion MRI and fMRI) in which all phase-encoding (PE) lines are read after a single radio frequency pulse. It was extended to single-shot EPI via gradient blips (Nunes et al., 2006), but this approach resulted in voxel tilting artifacts. Blipped-CAIPI acquisition uses sign and amplitude modulation of the gradient blips to avoid phase accumulation and eliminate the voxel tilting artifacts (Setsompop et al., 2012).

Consider an example from the HCP fMRI data, which uses AF = 8, FOV/3 blipped-CAIPI with left-right or right-left PE direction, an FOV of 208 × 180 × 144, and 2 mm isotropic voxels. The HCP acquisition protocol results in nine packets for each time point, where the number of slices in a packet is equal to the AF. The first packet contains slices 1, 10, 19, 28, 37, 46, 55, and 64. Without blipped-CAIPI, voxel [1,1,1] would be aliased with [1,1,10], [1,1,19], [1,1,28], [1,1,37], [1,1,46], [1,1,55], and [1,1,64]. With 90 voxels in the phase-encoding direction, phase shifts in k-space at increments of 2π/3 correspond to shifts in image space by 30 voxels, i.e., FOV/3. Then voxel [1,1,1] is aliased with [1,31,10], [1,61,19], [1,1,28], [1,31,37], [1,61,46], [1,1,55], [1,31,64]. Note some of the aliased locations are not shifted, here, [1,1,1], [1,1,28], and [1,1,55]. Variation in coil sensitivity along the z-axis is required to unalias these slices (Setsompop et al., 2012).

When a cluster of voxels share a common signal, the aliasing pattern can be calculated for all voxels in the cluster, which we call regional aliasing. Then signal may leak into the aliased regions and create clusters of false positives. A convenient approach to calculating aliased regions is to define a region of interest, simulate multislice acquisition from a mask of this region, and then reconstruct the image using either the original slice-GRAPPA kernel (hereafter, slice-GRAPPA) or split slice-GRAPPA. This approach provides insight into the magnitude of activation that results from slice leakage. We use a 5 × 5 kernel size (Uğurbil et al., 2013), which was used in the HCP reconstruction. The size of the kernel does not affect the locations of aliasing, although it could impact reconstruction accuracy and consequently the extent of leakage. A formal description of the slice-GRAPPA algorithm is in the Web Supplement; for a description of the split slice-GRAPPA, the reader is referred to [Cauley et al. (2014). To calculate aliased regions:

1. Create the real-valued input volume by defining the region (in the unregistered coordinate system of the scanner) where large activation is likely to occur. To define this region, one could estimate activation at each voxel, define a bounding box of the region of interest, and then threshold the test statistics in that region.
2. Simulate SMS acquisition:

a. Create complex-valued spatial-by-coil image data: multiply the real-valued input volume by the complex-valued coil sensitivities to create a four dimensional image. We estimated coil sensitivities as the ratio of the complex-valued channel data to the sum of squares of all channel data, which was then smoothed by a Gaussian filter with FWHM = 6 mm, and finally renormalized by the sum of squares of the smoothed data. As a consequence of this normalization, the sum of squares of the activation distributed across all channels is equal to the specified level of activation.
b. Create spatial-by-coil k-space data: apply the two-dimensional discrete Fourier transform (DFT) to the read-out and phase-encoding directions to convert data from the image domain to spatial frequencies.
c. Group the slices that will be aggregated into multislice packets: label the slices according to their multislice packet as in (S.1) in the Web Supplement.
d. Mimic blip-CAIPI acquisition: For each packet, apply phase shifts assuming perfect blip-CAIPI implementation. For FOV/3 shifts, this can be accomplished for the *c*th coil, *ℓ*th PE frequency, *k*th read-out (RO) frequency, and *z*th slice by 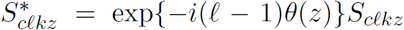 where 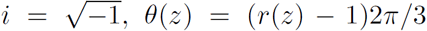 with *r*(*z*) ∈ {1,…, AF} indexing which slice *z* represents in the corresponding multislice packet and *S_cℓkz_* is the input volume from the previous step.
e. Generate the coil k-space multislice data: sum the slices in each packet as in (S.3).
3. Separate slices: create the output volume by applying the GRAPPA kernels as in (S.2), the 2D DFT, and sum-of-squares image reconstruction.

The reconstructed image can be compared to areas in which activation has been detected. An example of this procedure is depicted in Figure 1, which was created using the slice-GRAPPA kernel (left) and split slice-GRAPPA kernel (right) from the single-band reference data used in simulations. The input feature is a mask of the activation region used in the simulations in Section 3 multiplied by 100. The image reconstructed using this process contains non-zero values in the aliased locations, and the values represent the magnitude of leakage resulting from the input. In slice-GRAPPA, the greatest leakage occurs in an aliased region in brain tissue located inferior and left of the activated region. In the simulations in Section 3, we will see that this is an area in which we tend to detect slice leakage. In split slice-GRAPPA, the aliasing patterns are identical but the magnitude of leakage is greatly reduced. The statistical impact of slice leakage is also affected by the residual variances, which are impacted by noise amplification. Statistical impacts will be measured via specificity calculations in Section 3.

**Figure 1:**
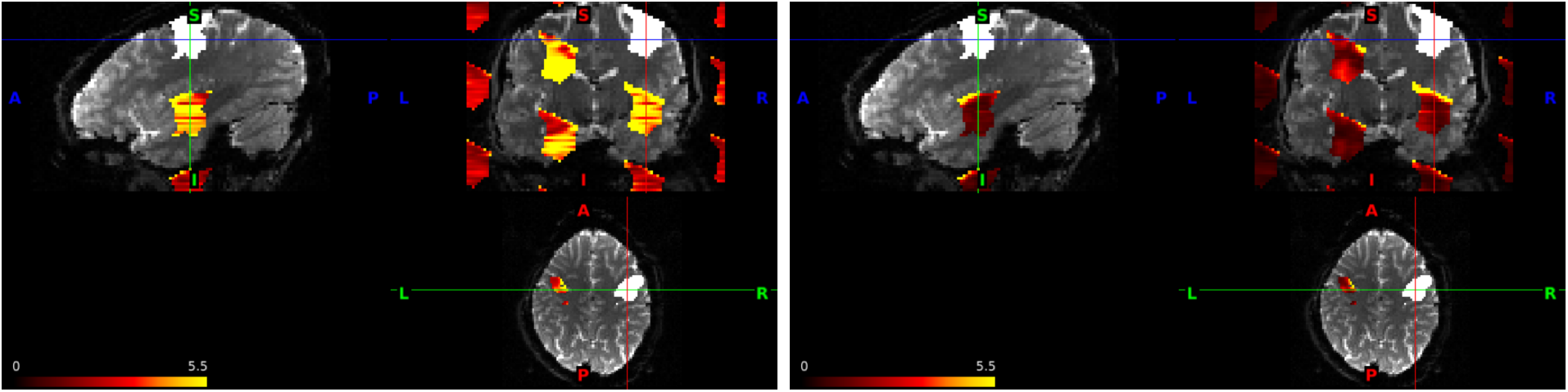
Activated voxels in simulations (white) and aliased locations for AF = 8 with FOV/3 shifts. The input volume is equal to 100 in the white region and zero everywhere else. Slice-GRAPPA (left) and split slice-GRAPPA (right), which have identical aliasing patterns but different magnitudes of leakage. In slice-GRAPPA, the output volume in the aliased regions has a maximum equal to 26.1, and in split slice-GRAPPA, the maximum is 25.6, but overall greatly reduced.

In practice, there is a stochastic component to whether and where slice leakage occurs. This will be seen in the simulations and in the examination of the unprocessed HCP data. Additionally, online scanner reconstruction algorithms may impact aliasing patterns. Vendor software used in the HCP includes corrections for spatial shifts due to respiration-induced B0 fluctuations (Uğurbil et al., 2013; Xu et al., 2013; Pfeuffer et al., 2002), which impacts the unprocessed HCP data. In another step, acquisition protocols may include online distortion correction due to gradient non-linearities (this is turned off in the HCP acquisition protocol and replaced by corrections during the preprocessing pipeline, Glasser et al. 2013, and thus does not affect the HCP unprocessed data).

## 3 Simulation Study

### 3.1 Simulation design

We conducted a simulation experiment with the following factors: acceleration factor (1, 4, 8); reconstruction method (slice-GRAPPA or split slice-GRAPPA); PE inter-slice image shifts corresponding to LR encoding (none or FOV/3); run duration (120 s, 240 s, 480 s, where the number of volumes depends on the acceleration factor); and activation magnitude (details below). We refer to the PE image shifts as CAIPI shifts as they simulate the effects of blipped-CAIPI EPI acquisition protocols. We simulated task fMRI data from a thirty-two channel head coil under conditions motivated by the HCP; see Section 4 for a description of the HCP unprocessed data. As a reference image, we used one volume of a navigator-corrected single-band fMRI of a subject collected during the pre-pilot phase of the HCP using their customized 3T Siemens scanner with a NOVA medical thirty-two channel head coil. The reference image contains 2 mm isotropic voxels with dimensions 104 × 90 × 72 in the read-out, phase-encoding, and slice directions.

The set of kernels for slice-GRAPPA reconstruction and split slice-GRAPPA was calculated from the calibration data for each level of AF × CAIPI shifts.

The initial design of the simulation study involved three steps:

1. Determine the scaling of the reference band image and the size of the measurement error to match empirical intensities and residual variances.
2. Determine the region of activation and baseline levels of activation.
3. Calculate activation time courses.

Step 1 is a challenging task because we do not have spatiotemporal-by-coil data available, but rather, magnitude data (i.e., the unprocessed HCP task fMRI data) and the single spatial-by-coil complex-valued reference image. We derived a measurement-error process in spatiotemporal-by-coil multislice k-space. The scaling of the magnitude image is complicated by the fact that taking the sum-of-squares reconstruction of mean-zero errors affects the mean structure. Concomitantly, the variance changes. The reference volume data intensities were in a different scale than the publicly available HCP single-band data. Consequently, we initially scaled it to have a mean equal to one and then determined the combination of mean intensity and residual variances to approximate the unprocessed fMRI data. Our strategy was to use the intensities in the non-tissue areas of the HCP unprocessed task fMRI data to inform the size of the variance of the k-space errors. Note the inverse discrete Fourier transform of independent and identically distributed (iid) Gaussian complex-valued errors with variance 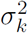 results in iid Gaussian errors but now the variances are scaled by the number of frequencies, 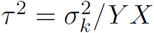, where *Y* is the number of frequencies in the PE direction and *X* is the number of frequencies in the RO direction. Then the square root of the sum of squares of mean zero complex-valued Gaussian noise is proportional to a chi random variable with 64 degrees of freedom (twice the number of coils due to the real and complex components), where the scaling factor is equal to τ. This is discussed in Aja-Fernández et al. (2011). The square root of the sum of squares of the Gaussian errors has a mean equal to 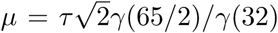 and variance 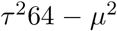, where γ(·) is the gamma function.

We chose to scale the reference volume in magnitude image space to result in a mean equal to 1,500. We added Gaussian complex-valued errors with both real and complex variance equal to 8.4573 × 10^7^ in k-space; in the absence of a mean signal, this results in a residual variance in magnitude image space equal to 4,500. This choice was informed by the empirical intensities and residual variances in non-tissue and tissue areas of single-subject massive univariate analyses of the HCP motor task data from the LR session of ten subjects. Our chosen parameters resulted in the median (across spatial locations) residual variance from the general linear model (GLM) fit to our simulated magnitude CAIPI AF = 8 data in non-tissue regions approximately equal to 3,700 (details of simulation below), where this value ranged from 1,800 – 4,300 in the ten HCP subjects. The median intensity in the non-tissue regions was 297 compared to 265–363 in the HCP data. Across all locations, the simulated median residual variance (with slice-GRAPPA reconstruction) was approximately equal to 6,700, whereas the HCP data ranged from 4,800–13,000, and for the median intensity, 450 versus 370–743. For the HCP data, these values were calculated from the white noise variance and intercept of the GLM described in Section 4.

For step 2, we constructed a single patch of activation based on an estimate of the region activated by the left-hand motor-task contrast from an HCP subject with a brain similarly shaped to our reference data. A GLM was fit to subject 101107 for smoothed and unsmoothed data from a motor task as described in Section 4. The smoothed data were used to roughly delineate the left-hand motor cortex, in which we defined a bounding box based on visual inspection, thresholded to voxels with approximate z-statistics greater than 2.326, and applied a brain mask. We then used the contrast levels from the unsmoothed data as the “true” activation, which contained some positive and some negative effect sizes (mean: 72.7, min: −293; max: 842, *n* = 1446). In our preliminary analysis, we set all voxels in the area of activation equal to the same level of activation, but this resulted in much larger t-statistics for the smoothed than unsmoothed estimates. Allowing the effect sizes to vary according to the unsmoothed contrasts resulted in more reasonable t-statistics. We then multiplied all voxels in the delineated region by a scaling factor to manipulate overall effect size.

For step 3, task onsets were simulated as a box-car function with the experimental condition lasting three seconds with onsets at 5, 65, 125, 185, 245, 305, 365, and 425 seconds for run duration = 480 s and the appropriate subset of times for run duration = 120 s and 240 s. The box-car function was convolved with the canonical hemodynamic response function from SPM (Penny et al., 2007). We performed simulations for scaling factors 0.5, 1, 2, 3, and 5. For scaling factor = 1, the average percent change relative to baseline was 1.06%, which was computed following Poldrack et al. (2011). Specifically, for each location in the region of activation, the true contrast value was multiplied by the peak of the task covariate, which equaled 0.48, then divided by the location’s mean from an fMRI times series simulated using the eight steps below with AF = 8, 240 seconds, and CAIPI shifts. Finally, we averaged this value across locations in the region of activation and multiplied by 100. For the other scaling factors, we estimated the average percent change as the corresponding scaling factor multiplied by 1.06%. Note that in the HCP design, the peak of the five motor task covariates (left hand, left foot, right hand, right foot, tongue) equaled 0.95, corresponding to 12 second tasks, while the peak from the cue was 3 seconds, with peak equal to 0.48; using the contrast from left hand versus others including cue (1 −1/5 −1/5 −1/5 −1/5 −1/5), multiplying by 0.95 and 100, and dividing by the mean, the average percent change from baseline from the subject used to define the region of activation (101107) was 1.61% from the subject’s unsmoothed data (in addition to the larger max of the task covariate, the mean fMRI time series for subject 101107 was somewhat higher than in the simulated data) and 1.21% in the smoothed data.

The simulation experiment then involved the following steps:

1. Create the (real-valued) activation time course for each voxel. For voxels in the activation region, multiply the baseline activation parameter by the scaling factor, and then multiply this result by the activation time course. All other voxels equal zero.
2. Create the true spatiotemporal image data by creating *n* copies of the (complex-valued) reference image, where *n* is the number of volumes, and then adding the activation time courses.
3. Create the true spatiotemporal-by-coil image data by multiplying each volume by the normalized complex-valued coil sensitivities (see step 2(a) in Section 2).
4. Group slices into multislice packets, convert the PE and RO directions to the frequency domain, and apply phase-shifts according to the CAIPI factor level (steps 2(b) - (d) in Section 2).
5. Add slices to create spatiotemporal-by-coil k-space multislice packets (step 2(e) in Section 2).
6. Add iid complex-valued measurement error to the spatiotemporal-by-coil k-space multislice packets assuming the same standard deviation for all spatial frequencies. This simple set-up produces realistic spatial variation in the residuals in the multiband data (Figure 2; for residuals based on HCP data, see Figure 7).
7. Use slice-GRAPPA or split slice-GRAPPA followed by sum-of-squares reconstruction to generate the simulated magnitude fMRI run.
8. Fit the GLM to each voxel. For this step, we conduct the analysis both with no smoothing and with smoothing using a Gaussian kernel with FWHM = 6 mm.

In these simulations, steps 1–5 are performed once for each level of (AF) × (CAIPI) × (run duration) × (activation magnitude). Then steps 6–8 are performed for each iteration. We conducted twenty-five iterations for each level. Note that in Step 6, we use iid errors across space and time. The iid errors in space allow us to determine that the resulting spatial variation in residual errors (Figure 2) is due to the SMS acquisition and coil geometry. With respect to time, simulating temporally correlated errors would decrease the effective sample size. We evaluate sensitivity and specificity with 120, 240, and 480 volumes, and thus cover a wide range of effective sample sizes. Results presented take approximately ten days on a 2.5 GHz Intel node with 256 gb memory and 48 logical processors.

**Figure 2:**
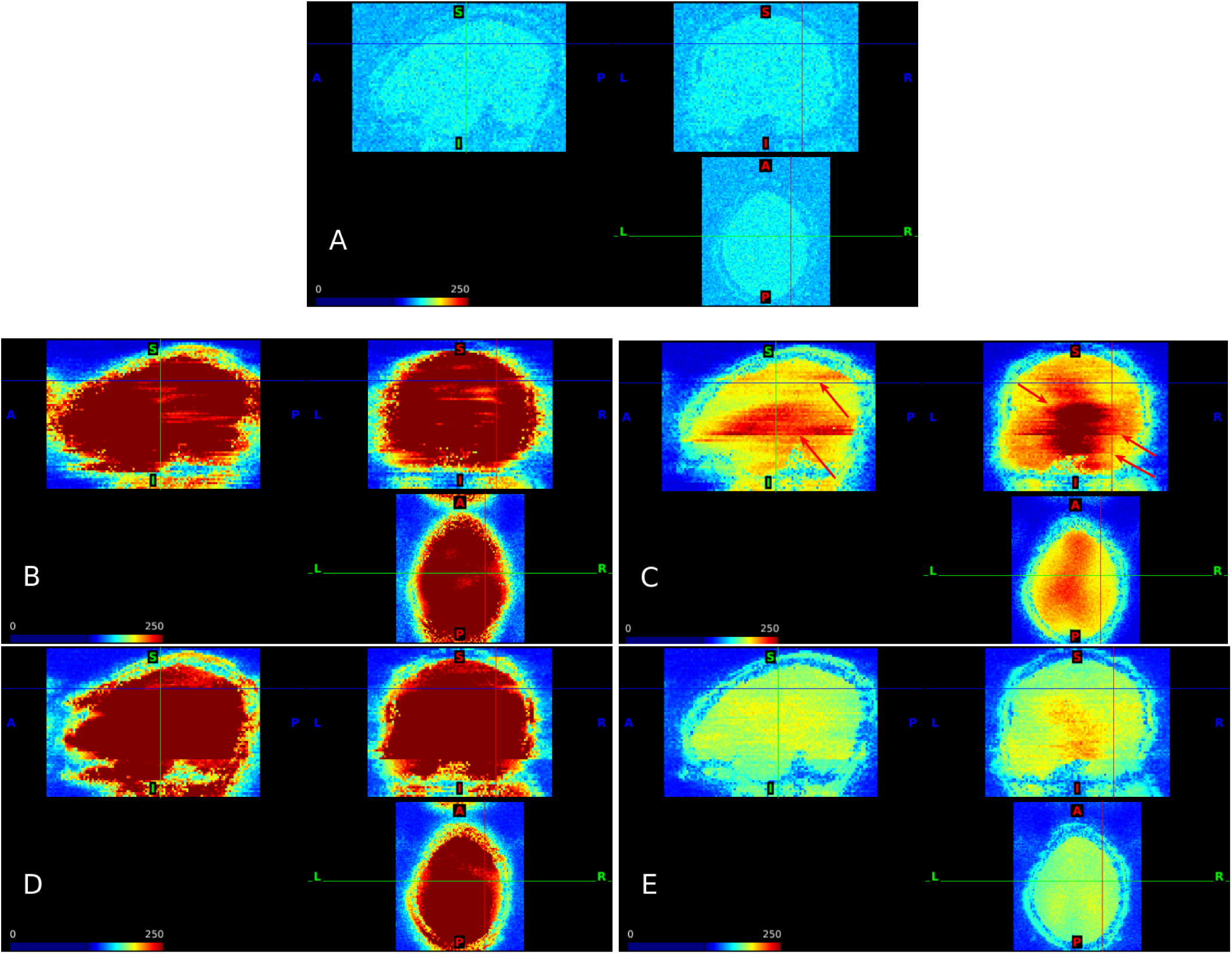
Noise amplification due to SMS using slice-GRAPPA reconstruction. Standard deviation of the residuals from the GLM fit to simulations with scaling factor = 1 and run duration = 480 s. AF = 1 (A); AF = 8 with no FOV shifts (B) and FOV/3 shifts (C); AF = 4 with no FOV shifts (D) and FOV/3 shifts (E). Results for split slice-GRAPPA and FOV/3 shifts are similar; see Web Supplement S.3.

We calculated sensitivity in each iteration based on the proportion of voxels with a p-value less than 0.001 among all true active voxels (*n* = 1446) from a two-sided t-test, and then averaged across iterations. We calculated one minus specificity in all locations, i.e., type one error rates, as the proportion of voxels among all inactive voxels (*n* =202,187) in the masked brain region with a p-value less than 0.001 from a two-sided t-test, and then averaged across iterations. We also calculated one minus specificity restricted to the set of voxels in the aliased regions intersected with the brain region.

### 3.2 Simulation results

#### 3.2.1 Noise amplification

The standard deviations of the residuals from the GLM fit to unsmoothed data provide insight into noise amplification due to SMS. We see that all SMS scenarios increase the standard deviation in brain regions relative to a single-band acquisition, but that FOV/3 shifts result in large decreases in the extent of noise amplification for both AF = 4 and AF = 8 (Figure 2). It is also apparent that there is a large cost to AF = 8 relative to AF = 4 for the case of FOV/3. The variance for AF = 4 and 8 is higher in subcortical and some prefrontal regions, particularly for AF = 8, which can be entirely attributed to SMS in our simulation design. We also see lines delineating higher and lower regions of variance in AF = 8, as highlighted by the red arrows in Figure 2. In the SMS literature, noise amplification is typically examined using *g*-factor maps, and our results are consistent with previous studies using *g*-factors (Todd et al., 2017; Setsompop et al., 2012). As described in Section 2, *g*-factors can be estimated as the ratio of the SMS standard deviations to the single-band standard deviations. These are provided in Web Supplement S.1. Results for split slice-GRAPPA and FOV/3 are very similar, while without FOV shifts split slice-GRAPPA tends to have somewhat larger standard deviations and *g*-factors; see Web Supplement Figures S.3 and S.4.

#### 3.2.2 Sensitivity and specificity

When the data were not smoothed, we found that sensitivity increased with AF when in conjunction with CAIPI but overall was low, while 1-specificity was generally maintained near the nominal alpha level for small to moderate effect sizes (Figure 3). When used with CAIPI, split slice-GRAPPA and slice-GRAPPA had very similar sensitivity, as indicated by the brown solid lines with filled squares (AF = S, slice-GRAPPA, FOV/S) versus brown dashed lines with hollow squares (AF = S, split slice-GRAPPA, FOV/S) in panels A-C, and similarly for AF = 4 with FOV/S (slice-GRAPPA in dark blue solid lines with filled triangles; splitslice-GRAPPA in dark blue dashed lines with hollow triangles). Without CAIPI, split slice-GRAPPA had somewhat reduced sensitivity relative to slice-GRAPPA (orange solid lines with filled squares correspond to AF = S, slice-GRAPPA, no shifts and orange dashed lines with hollow squares correspond to AF = S, split slice-GRAPPA, no shifts; light blue solid lines with filled triangles correspond to AF = 4, slice-GRAPPA, no shifts and light blue dashed lines with hollow triangles correspond to AF = 4, split slice-GRAPPA, no shifts; panels A-C). Split slice-GRAPPA improved specificity. In particular, the elevated 1-specificity in the slice-GRAPPA reconstruction represented by the brown solid lines with filled squares in panels D-I decrease to near 0.001 in most of the split slice-GRAPPA scenarios, represented by the brown dashed lines with hollow squares in panels D-I, which are difficult to see because they coincide with other scenarios. A zoomed-in view of specificity for the CAIPI acquisitions reveals split slice-GRAPPA resulted in good overall control of 1 - specificity (Figure S.5, A-C). Overall, multiband simulations with AF = S, CAIPI shifts, and split slice-GRAPPA reconstruction had better sensitivity with little slice leakage. Note that in these simulations, deviations of 1 - specificity from the nominal 0.001 level in Figure S D-F are due to slice leakage without the confounding effect of smoothing leakage.

**Figure 3:**
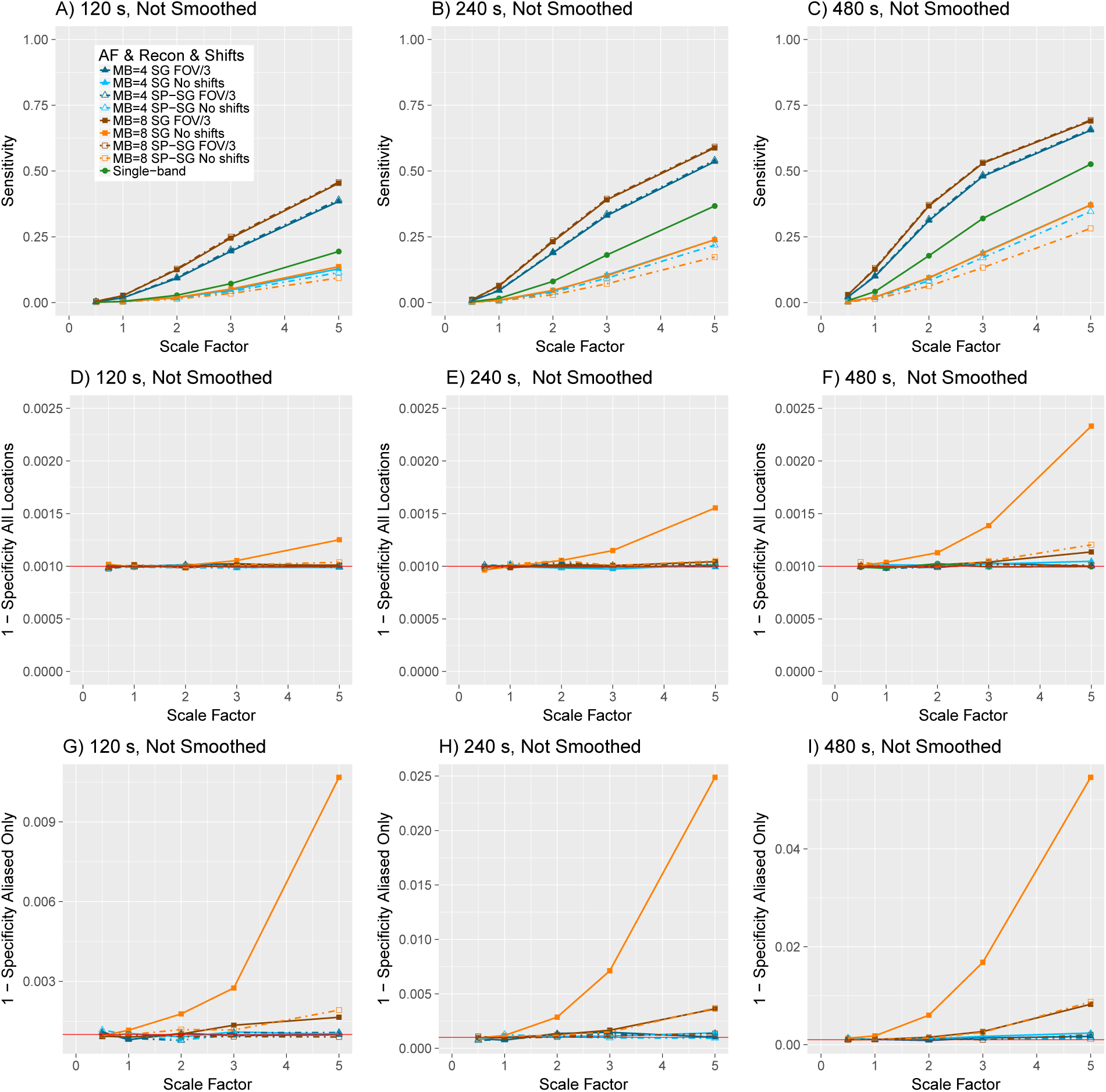
Sensitivity (power) (A-C), 1 - specificity (type 1 error rate) for all brain locations (D-F), and 1 - specificity (type 1 error rate) in aliased locations (G-I) in simulations with smoothing for *α* = 0.001. SG = slice-GRAPPA. SP-SG = split slice-GRAPPA. Scaling factors 0.5, 1, 2, 3, and 5 represent an average of 0.53%, 1.06%, 2.12%, 3.18%, and 5.30% change from baseline, respectively. See Web Supplement S.5 for a zoomed-in version of G-I.

When the data were smoothed, sensitivity increased with AF when in conjunction with CAIPI but at the cost of a decrease in specificity due to both smoothing leakage in voxels neighboring the region of activation and an increase in slice leakage (Figure 4). As before, split slice-GRAPPA and slice-GRAPPA had very similar sensitivity when in conjunction with CAIPI, as seen in the overlap in the brown solid lines with filled squares and the brown dashed lines with hollow squares (AF = S, slice-GRAPPA, FOV/S and AF = S, split slice-GRAPPA, FOV/3, respectively) and similarly, the overlap in the dark blue solid lines with filled triangles and dark blue dashed lines with hollow triangles (panels A-C). Without CAIPI, slice-GRAPPA tended to have higher sensitivity than split slice-GRAPPA, particularly for AF = 8 (orange solid lines with filled squares versus orange dashed lines with hollow squares, panels A-C). Over all brain locations, 1-specificity (panels D-F) was much larger than the nominal level for both reconstruction methods, where smoothing leakage impacts both. Restricting attention to the aliased regions, 1 - specificity for larger scaling factors is inflated for AF = 8 with slice-GRAPPA both with and without FOV/3 shifts (brown solid lines with filled squares and orange solid lines with filled squares) (G-I). Smoothing decreases the residual variances in the GLM, resulting in increased “power” to detect the slice leakage, such that the 1 - specificity rates are greatly increased in Figure 4 G-I relative to Figure 3 G-I. Split slice-GRAPPA reduces slice leakage, and most notably, generally controls 1 - specificity near nominal levels except for scaling factor = 5, as apparent in the zoomed-in depiction of the FOV/3 scenarios in Web Supplement Figure S.5, D-F. Thus the impacts on specificity in the split-slice GRAPPA simulations depicted in Figure 4 D-F are primarily driven by smoothing leakage to voxels neighboring the region of activation, and slice leakage is reduced to near nominal levels for small to moderate effect sizes (Figure 4 D-F, Figure S.5 D-F). The impacts of smoothing leakage and benefits of AF indicate practitioners can “tune down” the amount of smoothing used in preprocessing multiband data, as the boost in sample size from multiband can offset the loss in sensitivity from less smoothing.

**Figure 4:**
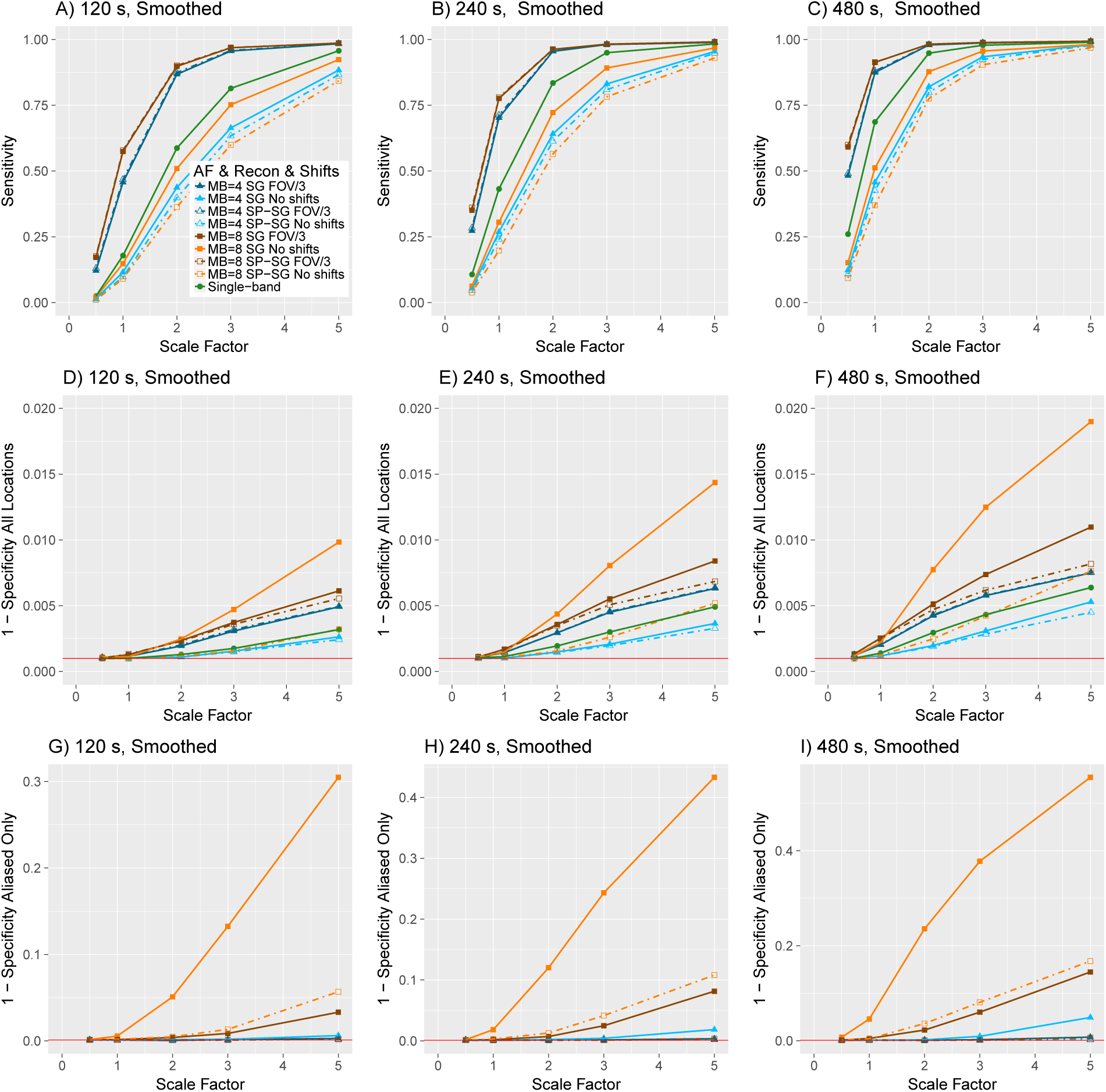
Sensitivity (power) (A-C), 1 - specificity (type 1 error rate) for all brain locations (D-F), and 1 - specificity (type 1 error rate) in aliased locations (G-I) in simulations with smoothing for *α* = 0.001. SG = slice-GRAPPA. SP-SG = split slice-GRAPPA. Scaling factors 0.5, 1, 2, 3, and 5 represent an average of 0.53%, 1.06%, 2.12%, 3.18%, and 5.30% change from baseline, respectively. See Web Supplement S.5 for a zoomed-in version of G-I.

CAIPI shifts always increased sensitivity, but the effects on overall specificity differed for AF = 4 and AF = 8 (Figure 4 D-F). AF = 4 with no shifts and split slice-GRAPPA had the lowest 1 - specificity (light blue dashed lines with hollow triangles, D-F), and AF = 4, no shifts, and slice-GRAPPA was similar (light blue solid lines with filled triangles, D–F). In the case of AF = 4 without CAIPI shifts and both slice-GRAPPA and split slice-GRAPPA reconstruction, the noise amplification is generally higher than AF = 8, no shifts, and slice-GRAPPA reconstruction, particularly in the core corresponding to the aliased locations (aliased regions fall in the same sagittal slices as the activation region), which can be seen in Web Supplement Figure S.2. This noise amplification also decreases the statistical impacts of leakage, leading to lower “power” to detect leakage. Consequently, AF = 4 has good specificity but at a large cost in terms of sensitivity, whereas AF = 8 with no CAIPI and slice-GRAPPA reconstruction has increased leakage as well as increased power to detect the leakage from both the lower noise amplification in the aliased regions and the increase in sample size from greater AF. Clearly, one would not want to use AF = 4 without CAIPI because of the large sensitivity costs.

These simulations also reveal that there can be a cost to split slice-GRAPPA in multiband acquisitions without CAIPI. AF = 8 with no shifts and split slice-GRAPPA reconstruction (orange dashed lines with hollow squares, Figure 4 A-C) has notably lower sensitivity than slice-GRAPPA (orange solid lines with filled squares). This corresponds to the increase in *g*- factor apparent by comparing the areas near the cross-hairs in the top-left panel of Figure S.1 to Figure S.4. This reveals that the impacts of CAIPI shifts, AF, and reconstruction method can be nuanced, although in all cases, CAIPI shifts improved sensitivity.

Inspecting the results from individual simulations provides insight into signal leakage. There appeared to be slice leakage in aliased regions in some simulations with AF = 8 and a more liberal alpha-level, which was reduced with AF = 4, and smoothing leakage was common (Figure 5). The variability suggests that slice leakage may be detectable in some, but not necessarily all, of the left-hand motor task contrasts in the unprocessed HCP data. Split slice-GRAPPA tended to reduce the size of the t-statistics in aliased regions, while maintaining the size of t-statistics in the region of true activation. For additional insight, in Section 3.3 we conduct simulations that isolate the effects of slice leakage from the features created by smoothing.

**Figure 5:**
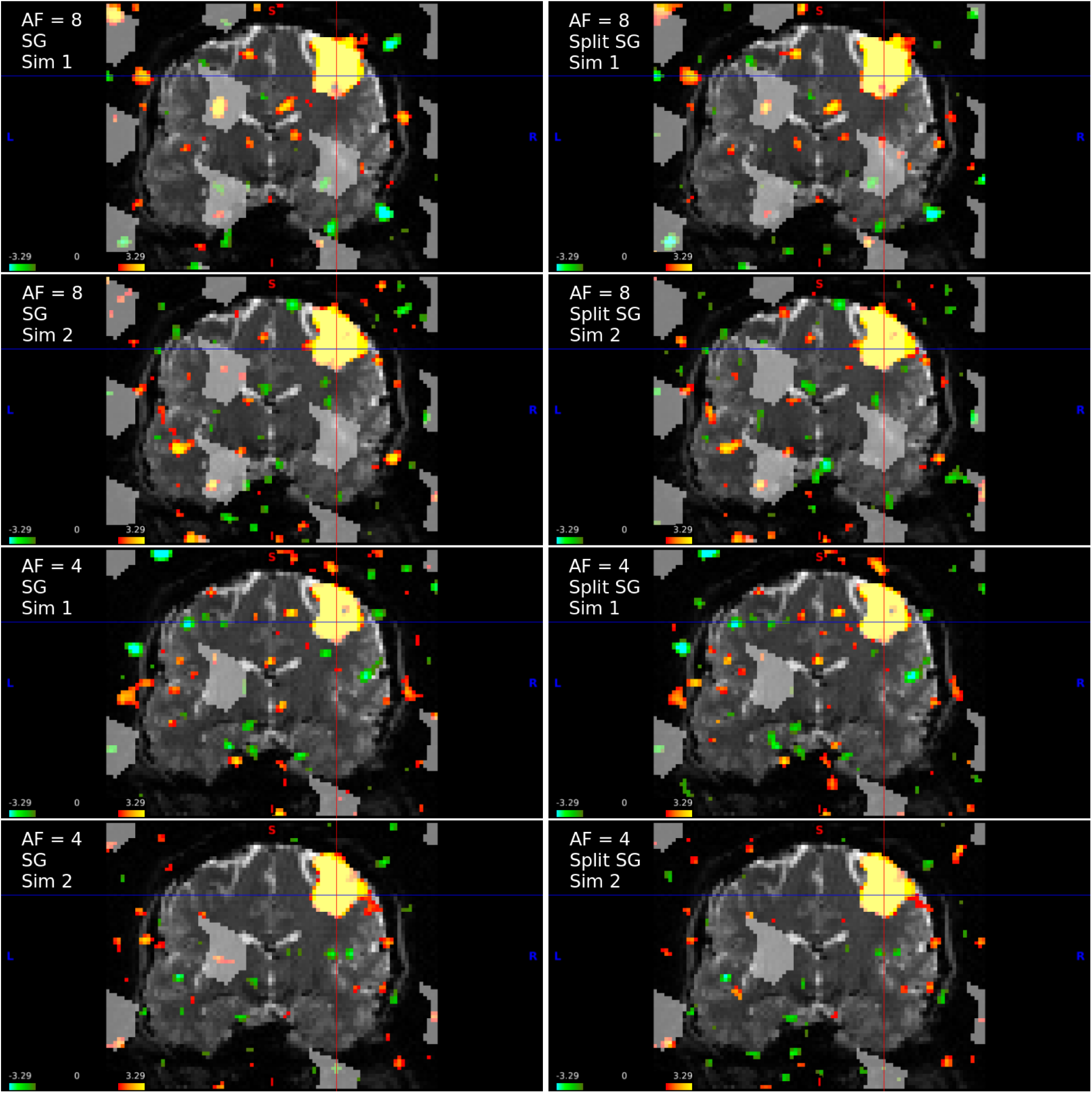
Example simulations for AF = 8 and AF = 4 for run duration = 240 s and FOV/3 shifts with scaling factor = 1 (average percent change from baseline = 1.06%). Shown are the t-statistics from two simulations for each AF. For AF = 8, the simulation depicted on the left shows some evidence of slice leakage (aliased locations in transparent gray) and the simulation on the right shows little evidence. Both simulations display evidence of smoothing leakage (the activated region leaking into neighboring voxels). Cursor at (63, 49, 53) in scanner coordinates. Thresholded at |*t*| > 2.

The benefits of split slice-GRAPPA are clearly visible at scaling factor = 5 and run duration = 480 s (Figure 6). For AF = 8, there is a dramatic reduction in slice leakage, and leakage is largely eliminated in the first simulation. Note that the leakage tends to be most prominent in the region in which the magnitude assessment of leakage from Section 2 was greatest (Figure 1). Simulations with AF = 4 have much less slice leakage for both slice-GRAPPA and split slice-GRAPPA reconstruction.

**Figure 6:**
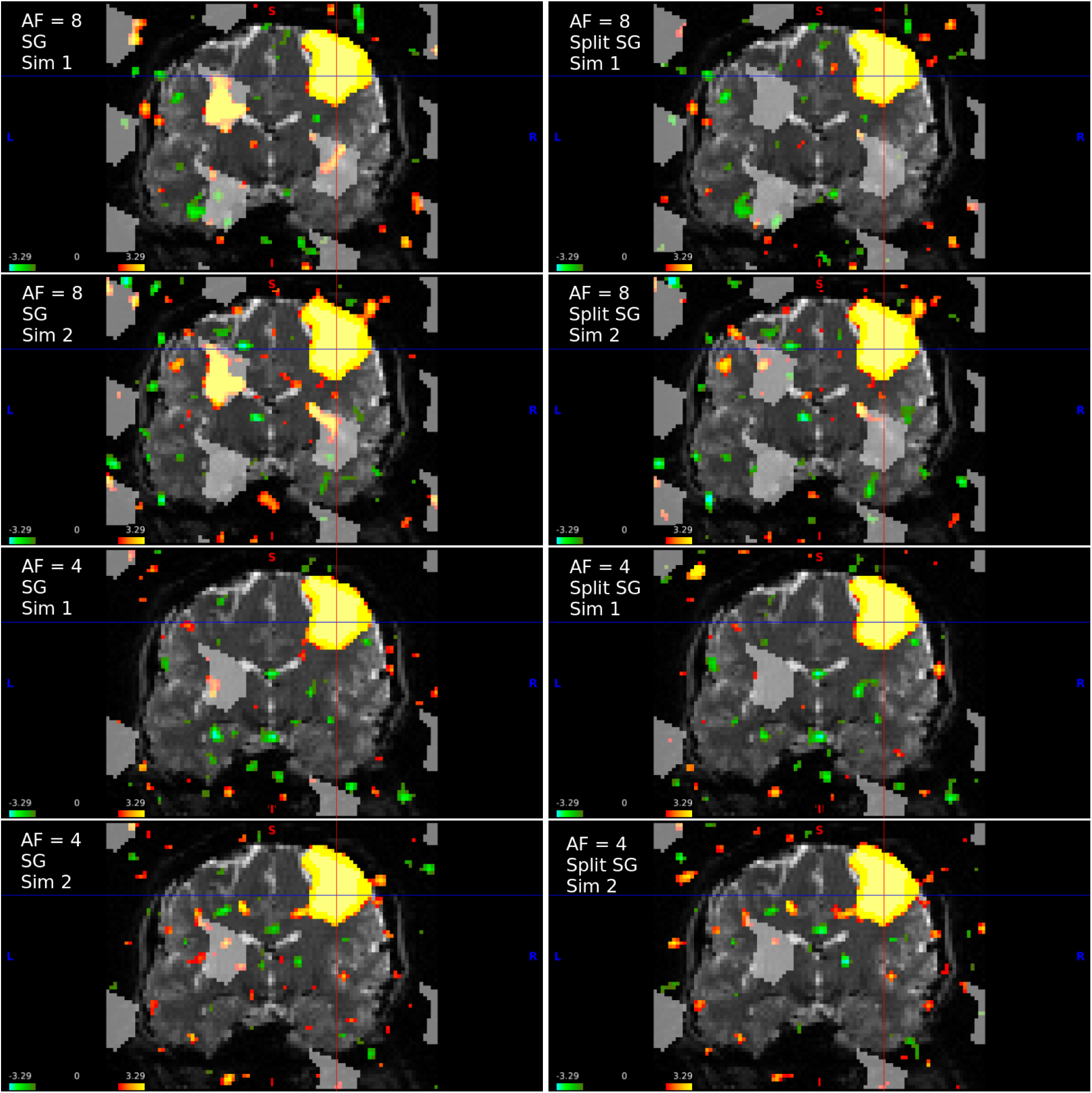
Example simulations for AF = 8 and AF = 4 for run duration = 480 s and FOV/3 shifts with scaling factor = 5 (average percent change from baseline = 5.30%). Shown are the t-statistics from two simulations for each AF. The left column shows slice-GRAPPA reconstruction and the right column shows split slice-GRAPPA reconstruction. Thresholded at |*t*| > 2.

### 3.3 Synthetic multiband simulation

To isolate the effects of slice leakage and the impacts of reconstruction, we first simulated two single-band datasets according to the design in the previous section: one for scaling factor = 1 and one for scaling factor = 5, where both datasets had run duration = 480 s, FOV/3 shifts, and TR = 1 s. We synthetically created multiband data from each single-band dataset by collapsing the single-band data into multiband packets and then applying slice-GRAPPA and split slice-GRAPPA reconstruction. In the previous section, the noise was added in k-space to the multiband packets, which produces *g*-factors found in previous literature (Figures S.1 and S.4; see for example Todd et al. (2017) for reference). In this section, the noise is added to the single-band data *before* collapsing into multiband packets. For the single-band data, this results in a realistic amount of noise. However, it introduces more noise to the multiband scenarios than the previous section, since for each 2D spatial frequency, a new noise term is generated for each slice. This contrasts with adding a single noise term to each 2D spatial frequency of the summed slices, as in Section 3.1. The alternative approach considered in this section produces unrealistically large *g*-factors in which areas susceptible to noise amplification have *g*-factors near eight rather than three for AF = 8. However, it has the advantage of allowing the same noise errors to be used in all scenarios, thereby allowing a direct comparison between the single-band ground truth and reconstruction method. We present results in which the TR = 1 s for all acquisitions alongside figures in which the data are subset to have TR = 2 s for AF = 4 and TR = 8 s for the single-band data (Web Supplement Figure S.6 and Figure S.7).

For scaling factor = 5, we see that AF = 8 has leakage when using slice-GRAPPA reconstruction. This is eliminated by split slice-GRAPPA in one aliased region, but a small cluster of leakage is still present in the region inferior to the region of activation (red arrows in Figure S.7). Note this cluster of activation is not present in the single-band data, and thus is attributed to slice leakage. Overall, there appears to be no cost to using split slice-GRAPPA instead of slice-GRAPPA for activation in the motor cortex according to this simulation design with FOV/3 shifts, as the regions of activation and t-statistics are very similar between the two reconstruction methods.

## 4 Slice leakage in the unprocessed HCP motor task data

### 4.1 Analysis of unprocessed HCP data

The HCP 3T motor task data contain two runs that differ in their phase-encoding direction. We searched for evidence of slice leakage by exploiting differences in the predicted aliasing patterns between RL and LR PE directions, as described below. HCP fMRI data use an AF = 8, blipped-CAIPI with FOV/3, no in-plane acceleration, FOV 208 mm × 180 mm × 144 mm, 0. 72 s TR, and 2 mm isotropic voxels (Smith et al., 2013), and each session of the motor task data contains 284 volumes. The acquisition parameters are summarized in Web Supplement Table S.1.

The unprocessed data represent the magnitude image space data that have been reconstructed online using the image reconstruction environment of Siemens scanners (ICE), which includes Nyquist-ghost correction, slice-GRAPPA reconstruction, and incorporates EPI intensity correction and corrections for shifts in space due to *B*_0_ off-resonance modulations. An example of the unprocessed data is in Web Supplement Figure S.4. The reconstruction algorithm was revised in April 2013. According to the HCP S900 Release Reference Manual, “the original reconstruction algorithm (version 177) performed the separation of the multi-band multi-slice images after transforming the acquired fully sampled data to frequency space along the read-out direction, and now the multi-band multi-slice separation is performed in k-space (WU-Minn HCP Consortium, 2015).” Both of the reconstruction versions used by the HCP are variants on the original slice-GRAPPA algorithm. Retrospective reconstruction using split slice-GRAPPA is not possible because the unreconstructed data were not saved due to storage limitations. Note our simulations perform separation entirely in k-space. Our use of the phrase “original slice-GRAPPA” refers to k-space reconstruction as described in Setsompop et al. (2012) and implemented in the second version of the slice-GRAPPA reconstruction algorithm used in the HCP, denoted r227. We used the ninety-eight subjects included in the U100 data sampler, which includes 57 subjects with r177 reconstruction, 39 with r227, and 2 with mixed. There was no significant effect of reconstruction version on the aliasing analysis, which is detailed below.

We used the unprocessed data because the locations of slice leakage are more predictable in the unprocessed data than any other stage of the preprocessing pipeline. There are a number of caveats to analyzing the unprocessed data. The HCP used a customized gradient that in particular improved diffusion imaging scans. Additionally, subjects’ heads were positioned off isocenter. These factors cause larger gradient nonlinearities than standard scanners. The fMRI volume pipeline of the HCP includes gradient distortion correction, motion correction to the single-band reference image, EPI image distortion correction to account for distortions due to RL and LR encoding, and non-linear MNI registration using FNIRT (Glasser et al., 2013). These processing steps change the aliasing patterns. Thus, we used the unprocessed HCP data. However, the unprocessed datasets do contain some preprocessing steps performed online during image reconstruction as previously described, and these could impact our ability to detect signal leakage.

For each subject, an AR(3) model with covariates was estimated for each voxel from the unprocessed data smoothed using a 6 mm FWHM Gaussian kernel, including non-brain voxels. The subject design matrices included the tasks convolved with the canonical HRF from SPM12 as well as the time-delay and dispersion derivatives for each task, a piecewise linear spline with four equally spaced knots to capture scanner drift, and the parameters from the affine registration used in the HCP preprocessing of these data. The voxel-wise AR(3) model can better account for increased temporal dependence from the shorter TR, spatially varying variances, and spatially varying temporal autocorrelation, and it was estimated using the reduced bias estimators from Worsley et al. (2002) and the code from the supplementary material in Risk et al. (2016).

To examine signal leakage, we defined a bounding box for the left-hand motor cortex equal to 51:78, 35:62, and 42:72 for the phase-encoding, read-out, and slice directions, which was based on inspecting the approximate z-statistics of the left-hand motor task contrasts from twenty subjects. For each subject, we created another map by thresholding the z-statistics from the left-hand contrast at 3.09. We then created a simple brain mask by thresholding the mean signal intensity across a given run. Lastly, we intersected these three maps.

Using the method described in Section 2, we created a mask of the regions aliased with the motor cortex from the LR run using the FOV/3 shifts, and we calculated “mismatched” aliased regions for the same run assuming the incorrect RL encoding, and vice versa for the RL run. An example of the aliased regions calculated according to the true PE direction and the aliased regions calculated assuming the incorrect PE direction is provided in Figure 8. Note that some regions with LR and RL encoding coincide. This is due to the cycling that occurs from using FOV/3 with AF = 8 and due to the chance occurrence that a voxel is aliased to some voxel in the activated region when using LR encoding and a different voxel elsewhere in the activated region when using RL encoding.

For each subject, we calculated the following quantities:

1. *A_LR–LR_:* Number of voxels with *z* > 1.96 in aliased regions of the LR run calculated using LR encoding.
2. *A_RL–RL_:* Number of voxels with *z* > 1.96 in aliased regions of the RL run calculated using RL encoding.
3. *A_LR–RL_:* Number of voxels with *z* > 1.96 in aliased regions of the LR run calculated using RL encoding.
4. *A_RL–LR_:* Number of voxels with *z* > 1.96 in aliased regions of the RL run calculated using LR encoding.
5. *N_LR_*: Number of voxels in aliased regions of the LR run (7*number of voxels in the activation mask).
6. *N_RL_:* Number of voxels in aliased regions of the RL run.

We then calculated the proportion of voxels activated in the “matched” aliased regions and in the “mismatched” aliased regions:

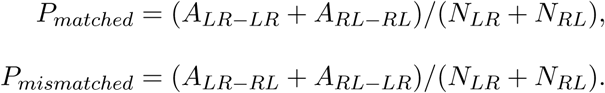

By combining the results from the RL and LR runs, this approach controls for possible confounding that would result if there happens to be greater true activation in the LR aliased regions, or vice versa. Activation in regions that are aliased in both matched and mismatched PE directions does not contribute to differences between *P_matched_* and *P_mismatched_;* these overlapping regions always occur in the unshifted aliased locations, and there is typically overlap in other regions. We designed our approach to ignore counts in these regions because slice leakage may be confounded with true activation and/or other artifacts. In this respect our approach is conservative.

To quantify evidence of slice leakage, we examined whether *P_matched_ > P_mismatched_* using a Wilcoxon signed-rank test. To check whether this analysis was affected by combining subjects between recon r177 and r227, we tested whether the difference in the proportion of voxels exceeding 1.96 in the matched and mismatched scenarios differed between r177 and r227 subjects, and there was no statistically significant effect (Wilcoxon rank-sum test: W=985, p=0.35), thereby justifying our simultaneous analysis of the ninety-eight subjects.

To examine SMS noise amplification, we fit the AR(3) models to the unsmoothed unprocessed data for four subjects (100408 LR, 103414 RL, 101309 LR, and 101915 RL), and examined the standard deviations. Specifically, we examined the estimated standard deviation of the white noise process from the AR(3) model, which is equal to the standard deviation of the residuals from ordinary least squares when the residuals are uncorrelated. (Note the first two subjects were reconstructed with r177 and second two with r227, and the noise amplification is apparent in all subjects.)

### 4.2 Results

#### 4.2.1 SMS noise amplification in unprocessed HCP task data

Our analysis of the unsmoothed HCP data revealed noise amplification from SMS acquisition. Similar to Section 3.2.1, there is a consistent pattern of higher variance in subcortical areas (Figure 7). Additionally, noise amplification is apparent in prefrontal regions. There were boundaries between axial planes that separated regions of higher and lower variance, which are particularly evident when viewing slices in the middle of the brain (right column in Figure 7). Boundaries in the *g*-factor are also apparent in the sagittal view in Figure 2 for AF = 8 in Todd et al. (2017). For studies mapping activation in a feature that extends across these boundaries, the activation maps would be spatially “biased” by differences in sensitivity, in particular delineating the feature in the low-variance region but potentially missing it in the higher variance region. The motor cortex does not appear to straddle such high and low noise amplification regions, but this may be an important issue for other tasks.

**Figure 7:**
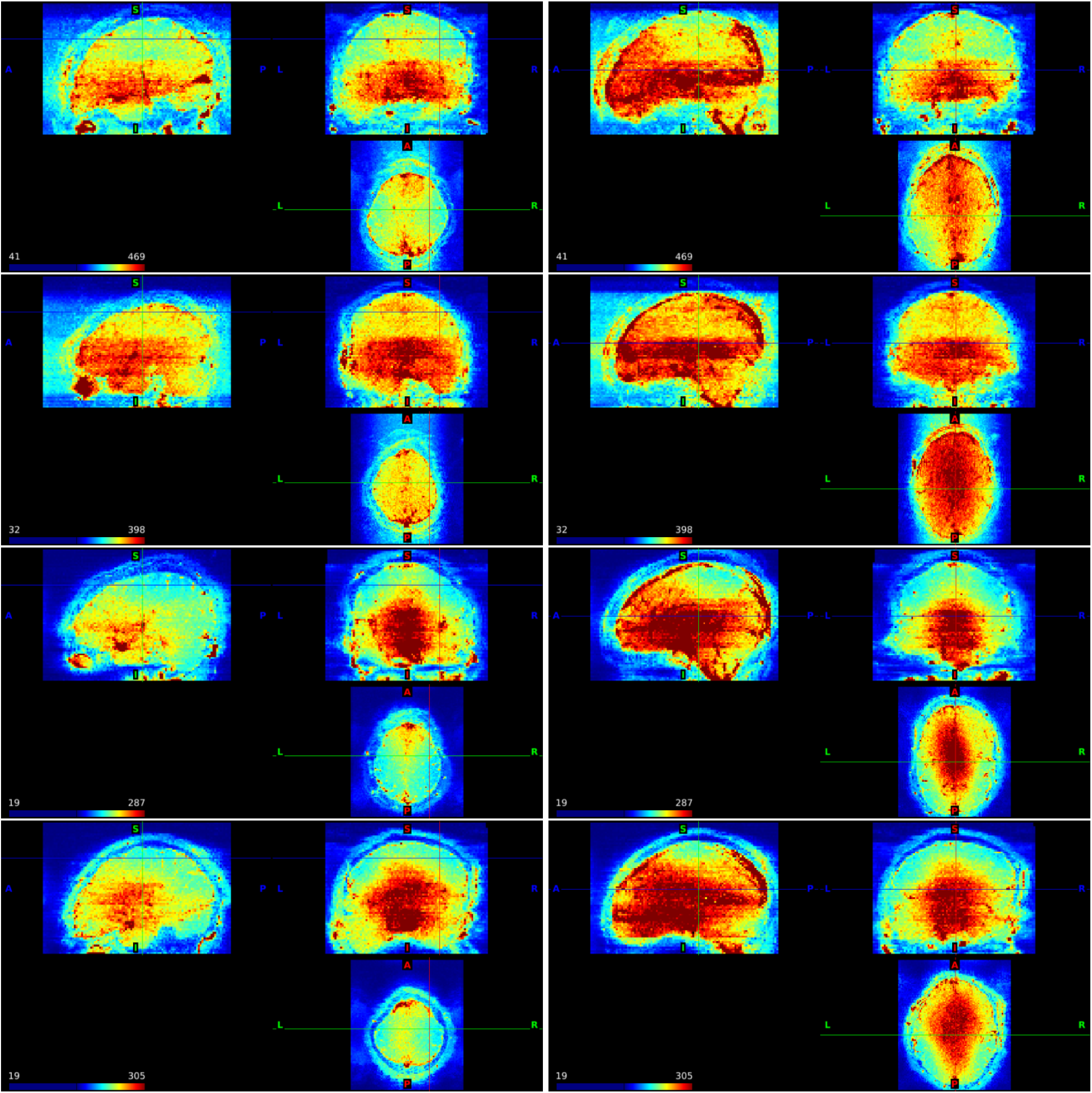
The standard deviation of the residuals from the point-wise AR(3) models fit to the unprocessed HCP motor task data from four subjects with the cursor at (63, 49, 53) (left column) and (46, 44, 37) (right column) in scanner coordinates. First row: 100408 with RL encoding; second row: 103414 LR; third row: 101309 LR; and fourth row: 101915 RL.

#### 4.2.2 Slice Leakage

Overall, we found evidence of slice leakage in many, but not all, subjects. The proportion of voxels with *z >* 1.96 was significantly greater in the matched than mismatched aliased regions (p<1e-09, Table 1), with 81 of 98 subjects having higher counts in their matched regions. Detailed counts are provided in Web Supplement Tables S.2 and S.3. To gain insight into our methodology, Figure 8 shows an example of the LR run from a subject in which voxels in LR aliased regions tended to have larger approximate z-statistics than voxels in the RL aliased regions. In this figure, the red arrow in the coronal slice indicates voxels with large z-statistics that are aliased with LR but not RL phase encoding. There are many other clusters with z-statistics greater than 1.96. Note we do *not* assess whether leakage is significant in any individual subject. Our methodology is unaffected by issues of multiple testing because we look at the counts of voxels in the matched versus mismatched regions, rather than make inference at the voxel-level, and thus we account for random clusters related to smoothness. It also accounts for regions of true activation and/or other artifacts by combining the RL and LR runs.

**Figure 8:**
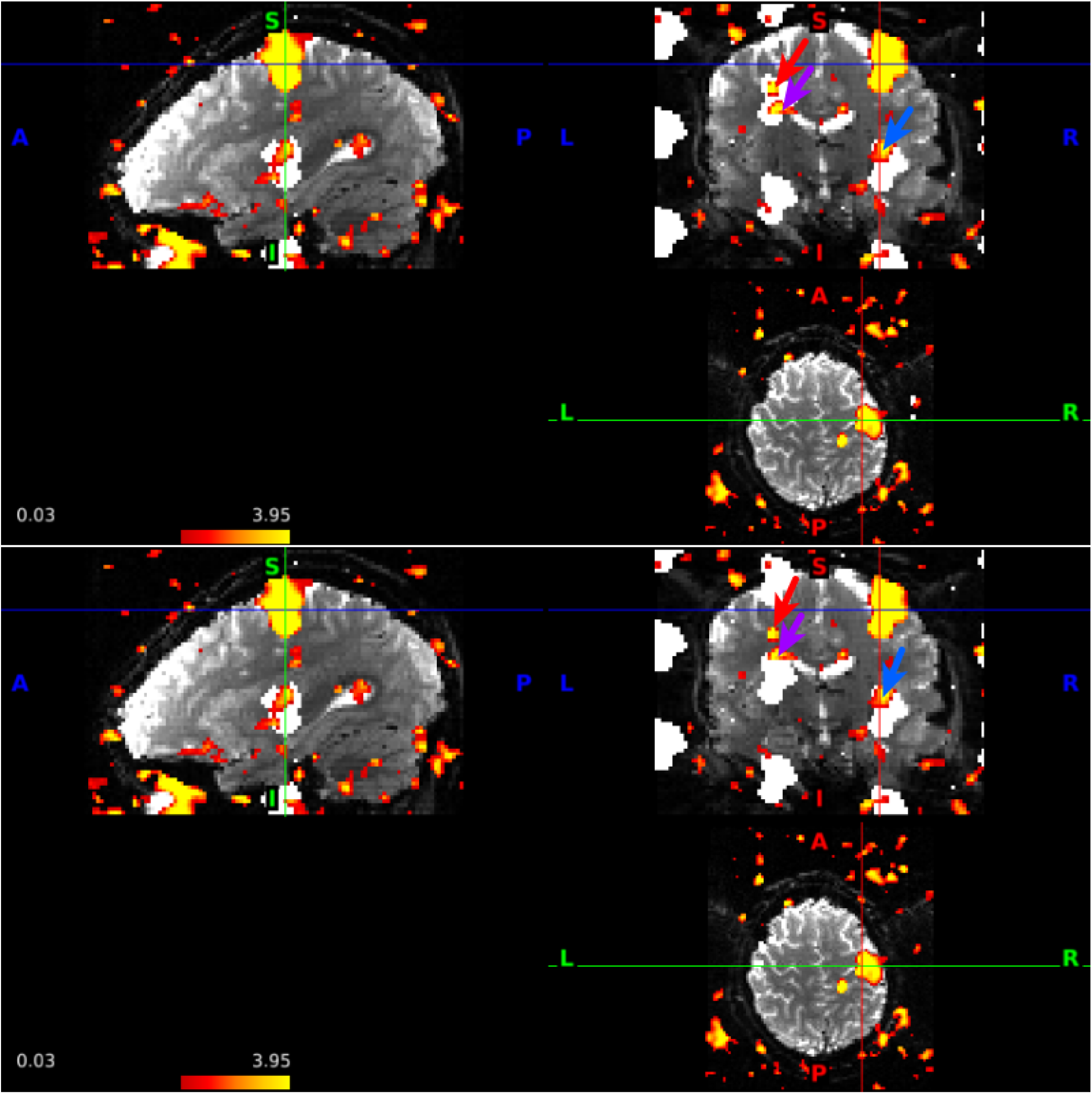
An illustrative example of voxels that contribute to evidence of slice leakage using our methodology (red arrow) in the LR run of subject 103111. Significance is not assessed at the subject or voxel level, but rather across all subjects. The regions aliased in the LR PE run are depicted in white in the top figure. The red arrow indicates a cluster of voxels with *z* > 1.96 that are aliased to voxels in the left-hand motor cortex using LR PE but not the RL PE. The bottom figure depicts the aliased regions that would result with RL PE, which differ in the coronal slice. The purple arrow points to a cluster in which some voxels fall exclusively in the LR aliased region and thus contribute to evidence of slice leakage, while other voxels fall in the region of overlap between the LR and RL regions and do not contribute to evidence of slice leakage. The blue arrow points to voxels aliased in both LR and RL PE directions, but do not contribute to evidence of slice leakage in our methodology.

**Table 1:**
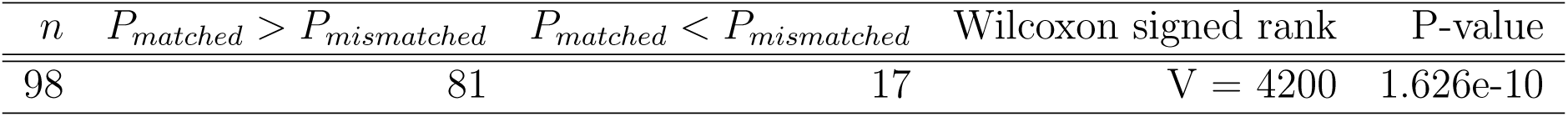
Number of subjects from the unrelated 100 HCP release in which the proportion of voxels with *z* > 1.96 was greater in the predicted aliased regions (matched) than in regions that are aliased using the incorrect PE direction (mismatched). A larger proportion in *P_matched_* is considered evidence of slice leakage. Significance assessed using a one-sided Wilcoxon signed rank test.

| $n$ | $P_{matched} > P_{mismatched}$ | $P_{matched} < P_{mismatched}$ | Wilcoxon signed rank | P-value |
| --- | --- | --- | --- | --- |
| 98 | 81 | 17 | V = 4200 | 1.626e-10 |

Our retrospective examination of slice leakage in the unprocessed HCP data is limited to examining locations that differ between the RL and LR runs, and this limitation eliminates some of the aliased regions because they are equivalent in the matched and mismatched PE directions. It is also unable to distinguish slice leakage when portions of aliased regions that are affected by shifts happen to overlap, which is common when the extent of activation is large. A cluster of higher-valued z-statistics was found in the aliasing region inferior to the motor cortex in many of the subjects, but since the region is aliased in both the LR and RL directions, activation in this region in our approach is considered equivocal. The blue arrow points to one of these regions in Figure 8. Figure 9 depicts the LR run from another subject, and the lower blue arrow is located 27 slices away from the blue arrow in the left-hand motor cortex. The brown arrows in the coronal slice depict two clusters that are located 27 slices apart and aliased with each other, but are not quite aliased with the left-hand motor cortex, and thus also do not contribute to evidence of slice leakage in our analysis. If a group analysis of HCP data were conducted, sensitivity would be increased by combining the two runs from a subject, but when aliased regions coincide in both runs (e.g., the blue arrows), there is an increased ability to incorrectly identify these regions as task activated. In an examination of the publicly available HCP analysis of the preprocessed data for subject 101915 combining the LR and RL runs, the region of activation identified by the lower blue arrow of Figure 9 approximately corresponds to MNI coordinates (40,-16,-14); in a map formed by the keyword “motor cortex” in NeuroSynth (Yarkoni et al., 2011), this location is not associated with motor tasks.

**Figure 9:**
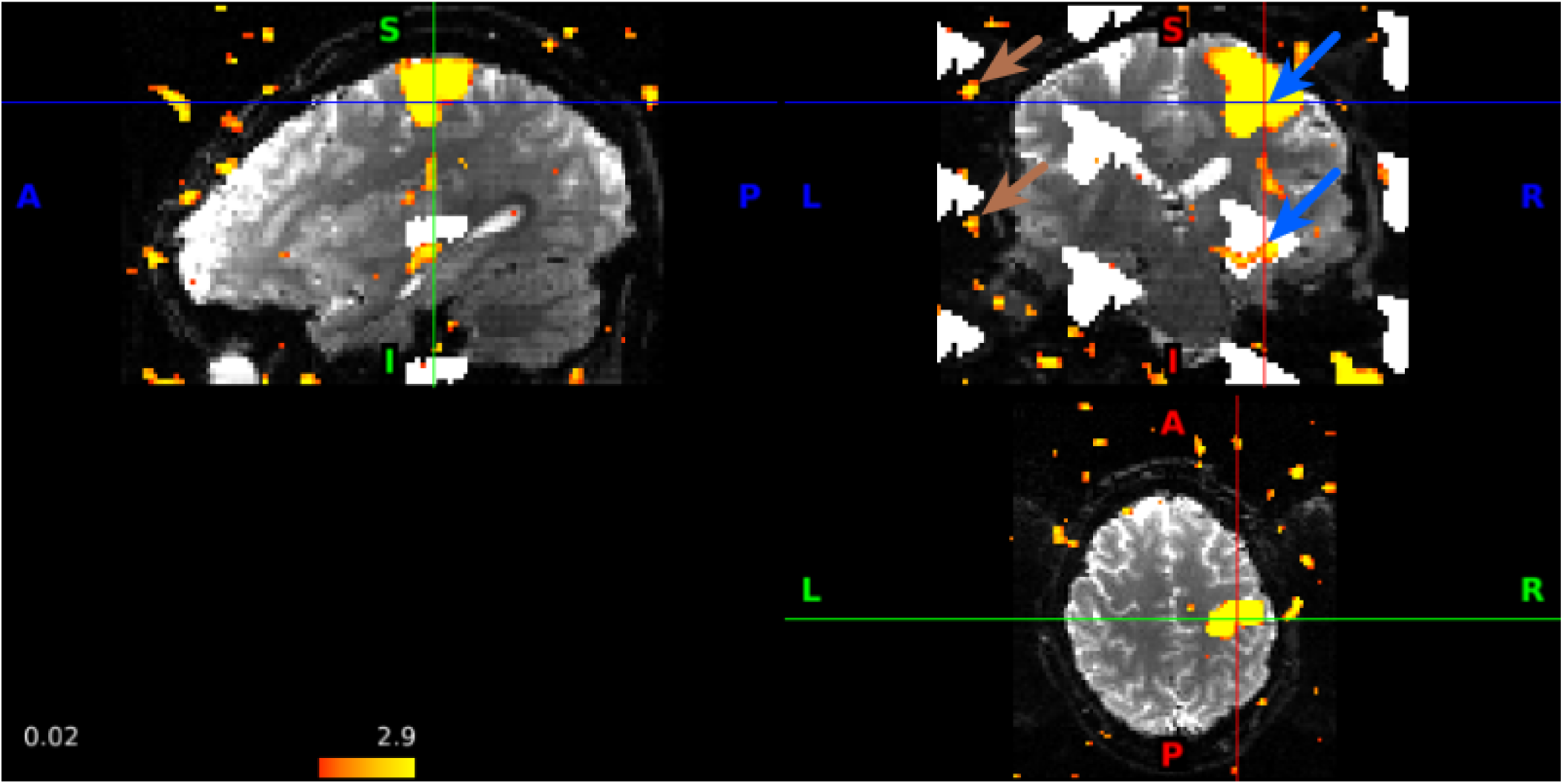
Equivocal evidence of slice leakage in the LR run of subject 101915 that is not attributed to slice leakage in our methodology. The blue arrows in the sagittal and coronal views are located at (63,44,54) and (63,44,27) in the scanner coordinate system, and these locations are aliased in the AF = 8 and FOV/3 acquisition protocol for both LR and RL encoding. The brown arrows in the coronal view are examples of aliased locations (7,44,57) and (7,44,30) in which the source of the signal is uncertain; they are close to but not contained in the LR aliased regions of the motor cortex (white) (nor are they contained in RL aliased locations). Thresholded at *z* > 1.96.

There is a large stochastic component to the extent of slice leakage and whether we are able to detect it at the effect sizes observed in the single-subject analyses. To gain insight into our leakage detection approach, we conducted 100 simulations according to the design in Section 3 using run duration = 240 s, AF = 8, CAIPI shifts, scaling factor = 1, and slice-GRAPPA reconstruction. In these simulations, we found that *P_matched_ > P_mismatched_* in 76 of the simulations, and *P_matched_* was significantly greater than *P_mismatched_* (V = 4331, p < 0.0001). We subsequently conducted the same analysis with scaling factor = 5, and *P_matched_ > P_mismatched_* in 100% of simulations.

We then conducted the same simulation experiment using split slice-GRAPPA reconstruction. For scaling factor = 1, *P_matched_ > P_mismatched_* in 56 / 100 simulations, and *P_matched_* was not significantly greater than *P_mismatched_* (V = 2863, p = 0.12). For scaling factor = 5, *P_matched_ > P_mismatched_* in 93 simulations (V = 4956, p < 0.0001).

Note the visual examination of the results from our simulation study also found a stochastic component to whether leakage was detected, even when we were not limited to examining differences in regional aliasing patterns due to PE direction (Figure 5). This is analogous to a low-powered (less sensitive) experiment in which a signal is not always detected, since we know there is a true signal from leakage albeit an undesired one. Thus, small effect sizes have less risk of detectable leakage.

## 5 Exploration of ICA-FIX preprocessed HCP resting-state data

An important question is whether SMS impacts persist in the processed data. The HCP has released resting-state data in which artifacts were identified using ICA and their variation subtracted from the resting-state time series, as described in Salimi-Khorshidi et al. (2014) and Smith et al. (2013). In some instances, ICA can identify artifacts that are related to SMS acquisition. An example is in Figure 8 in | Griffanti et al. (2016), which is described as a “checkerboard” effect corresponding to activation every eighth slice for AF = 8 and an acquisition with 64 slices, which appears to correspond to head motion. We speculate that the banding pattern may be related to subject movement along the slice direction, leading to reconstruction using the GRAPPA-kernel trained on a different part of the brain.

To gain insight into noise amplification from SMS in the preprocessed data, we examined the standard deviation of the fMRI time courses of the ICA-FIX resting-state data in volume space. Figure 10 presents the standard deviations for the following datasets: rfMRI_REST1_LR_hp2000_clean.nii.gz from subject 100408 (r177 reconstruction); rfMRI_REST2_LR_hp2000_clean from 103414 (r177 reconstruction); rfMRI_REST1_RL_hp2000_clean.nii.gz from 101309 (r227 reconstruction); and rfMRI_REST2_RL_hp2000_clean.nii.gz from 101915 (r227 reconstruction).

**Figure 10:**
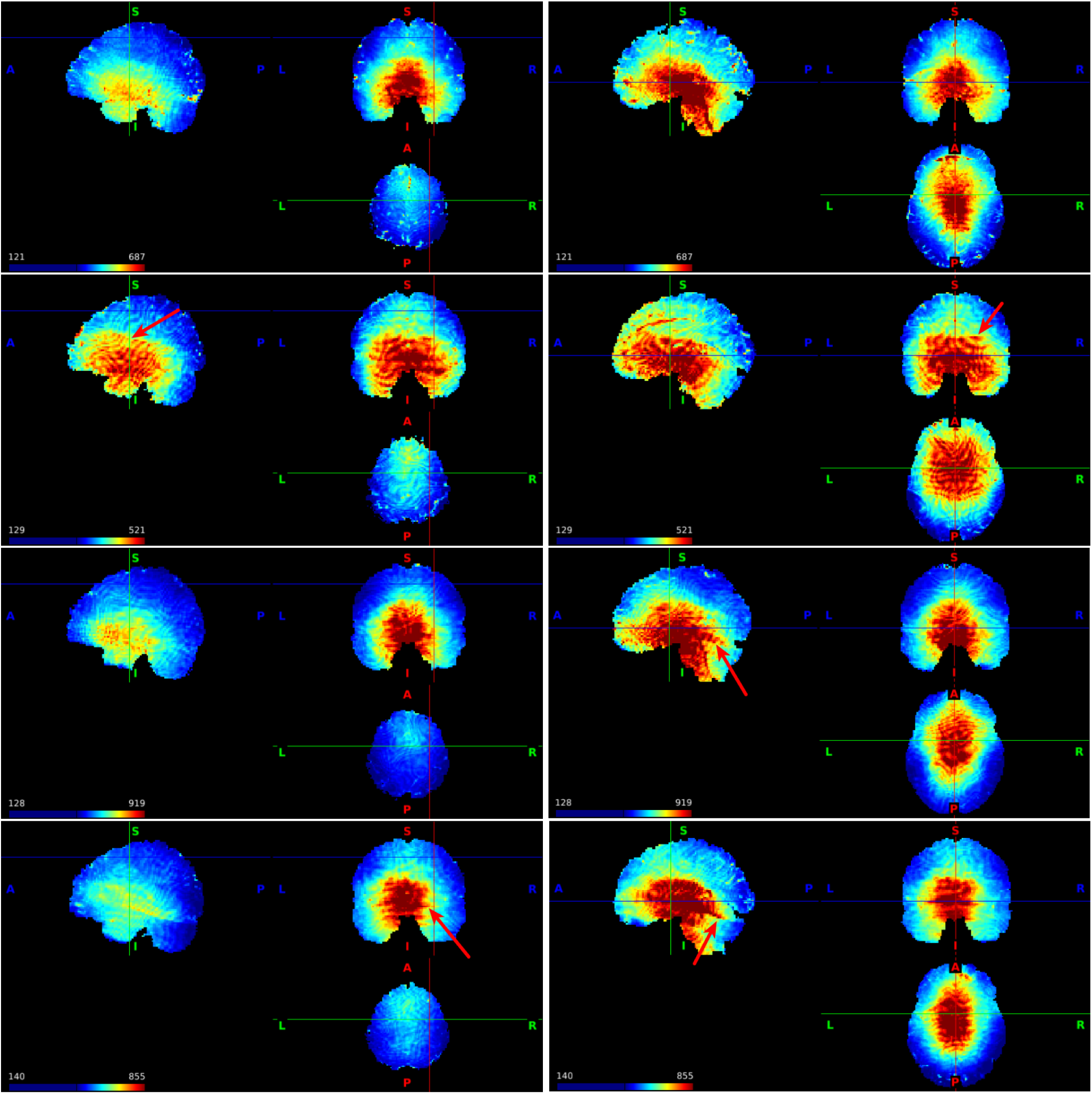
Standard deviations of the time series for the preprocessed, ICA-FIX resting-state data from four subjects with the cursor at (32, −8, 54) (left column) and (0, 0, 0) (right column) for subjects 100408 (first row), 103414 (second row), 101309 (third row), and 101915 (fourth row).

The standard deviations are notably higher in subcortical regions in all subjects (Figure 10). This may be due to multiple factors including noise amplification from SMS acquisition, as occurred in our simulations (Figure 2). Note that when 32 or 64-channel head coil arrays are used, the signal in subcortical regions can be attenuated, and normalizing the signal can also contribute to increased standard deviations. In some instances, boundaries between higher and lower variances due to SMS acquisition are apparent, where the lines that were seen in the task standard deviations in Figure 7 are now rotated approximately 20 degrees in the sagittal views, corresponding to registration of the oblique slices. Overall, the boundaries are blurred, which may be related to a variety of factors including motion correction, ICA-FIX, and the interpolation steps during preprocessing and registration that result in a small degree of smoothing.

## 6 Discussion

There are many benefits of SMS techniques, which can lead to novel scientific insight. However, we suggest that there are important costs in terms of structured noise amplification and specificity. Our findings indicate that slice leakage may have impacted the results of previous studies that used the original slice-GRAPPA (no leak block) reconstruction, and further, that future studies using publicly available datasets acquired with the original slice-GRAPPA reconstruction should be aware of these impacts when interpreting task activation. Our simulations indicate specificity can be improved by using split slice-GRAPPA (leak block) and less smoothing. Structured amplification of measurement error persists in split slice-GRAPPA but can be attenuated by moderate acceleration factors (AF = 4). Our simulation study complements the results of previous empirical studies that compare test statistics when the truth is not known, where it is difficult to attribute whether an increase in the size of test statistics and/or the number of activated voxels is due to improvements in sensitivity or decreases in specificity. We disentangle the effects into true and false positives. We found evidence of slice leakage in smoothed, unprocessed motor task data from the HCP, which used the original slice-GRAPPA reconstruction. We also found evidence of noise amplification due to SMS in both unprocessed motor task data and the ICA-FIX resting-state data.

We focused on a case study of activation in the left-hand motor cortex using coil sensitivities from the HCP Siemens Skyra 3T scanner with 32-channel head coil. Our simulations indicate AF = 8 may be preferred for mapping motor cortex, in which slice leakage can be reduced by split slice-GRAPPA and overall specificity can be improved by decreasing smoothing. The slice leakage at AF = 8 was substantial with slice-GRAPPA in smoothed data but was reduced by split slice-GRAPPA to near nominal levels for all but the largest effect sizes (Web Supplement Figure S.5 D-F). The simulation results for smoothed data in Figure 4 indicate SMS with split slice-GRAPPA has more smoothing leakage than single-band acquisitions, but a more useful way of looking at this result is that the amount of smoothing can be reduced in SMS acquisitions because the boost in sensitivity from the acquisition of more volumes helps offset the sensitivity cost from decreasing smoothing. Thus, SMS offers a means of reducing smoothing leakage by facilitating the use of a smaller FWHM.

Sensitivity is one way to measure whether the boost in effective sample size in SMS outweighs the cost of noise amplification, but our examination of the spatial patterns in noise amplification suggest other potential costs that are decreased by moderate acceleration factors (AF = 4). Figures 2, 7, and 10 indicate possible costs not captured by our sensitivity simulations. First, the boundaries between high and low variance could create biased estimates of spatial activation patterns when a cluster of activation spans high and low variance regions. Secondly, we simulated activation in the motor cortex, and subcortical and prefrontal regions may have optimal sensitivity at lower AFs. The left-hand motor cortex used in our study represents a low *g*-factor area. Todd et al. (2017) found that the benefits of SMS were higher in the low *g*-factor noise area of V1, moderate in the moderate *g*-factor area in the parahippocampal place area, and sometimes decreased t-statistics in the ventral-medial prefrontal cortex. Their results indicate that our estimates of sensitivity and specificity are specific to our choice of activation area. For studies targeting subcortical and some prefrontal regions, the standard deviation maps in Figures 2 and 7 and results from Todd et al. (2017) suggest a lower multiband factor may be preferred for improving sensitivity, which is independent of leakage considerations. This would have the dual benefit of also decreasing leakage. In addition to the spatial impacts, the costs and benefits of SMS may depend on factors such as scanner, head coil, and phase encoding direction. | Demetriou et al. (2016) found that the benefits of SMS did not differ greatly between scanner platforms but were affected by the particulars of the analysis method, e.g., voxel-based, ROI-based, or multi-voxel pattern analysis, where benefits were found in task fMRI using MVPA but not the ROI-based approach.

Our motor task simulations indicate split slice-GRAPPA improves specificity over slice-GRAPPA without a decrease in sensitivity when using FOV/3 shifts, although issues with structured noise amplification in AF = 8 persist. Cauley et al. (2014) notes that split slice-GRAPPA can increase the reconstruction error for the fMRI time series, but it may be more robust in diffusion weighted imaging where the contrast varies substantially from the training data. Todd et al. (2016) found significant differences in temporal SNR between slice-GRAPPA and split slice-GRAPPA, but did not find significant differences in the number of activated voxels. In our simulations, split slice-GRAPPA with FOV/3 shifts did not have higher noise amplification than slice-GRAPPA (Web Supplement Figures S.3 and S.4, Figure |2). The results suggest split slice-GRAPPA should always be preferred to slice-GRAPPA in HCP-like acquisitions with LR phase encoding and FOV/3 shifts. We caution that the impacts may differ in acquisitions using anterior-posterior (rather than LR) phase-encoding, and that these findings are sensitive to the choice of FOV shifts. In particular, the simulations showed that there was a cost in terms of sensitivity to split slice-GRAPPA without FOV/3 shifts (Figure 4), which raises the possibility that ideal FOV shifts for maximizing sensitivity in slice-GRAPPA may differ from split slice-GRAPPA. Assessing these trade offs is an important avenue for future research.

Our simulations demonstrate large decreases in noise amplification and associated improvements in sensitivity when using FOV/3 shifts relative to no shifts. Utilizing the same FOV/3 shifts across different AFs may not be optimal. Blipped-CAIPI protocols to achieve other FOV shifts may decrease noise amplification. Careful choice of shifts and AF could be designed to decrease the risk that regions aliased with a focal region are located in the brain and/or gray matter, such that the shifts are chosen to relegate aliased regions to areas of the image space that are masked in a statistical analysis. The development of 3D acquisition protocols with corkscrew trajectories (Bilgic et al., 2015) or the use of “incoherent” FOV shifting (Zhu et al., 2014) may diminish the patterns of noise amplification apparent in Figure 7.

An interesting question is how the noise amplification impacts functional connectivity studies. An increase in measurement error leads to a decrease in correlation. Thus, high AF may lead to lower functional connectivity for edges in which one or both locations are in areas with a high *g*-factor. We suggest that moderate multiband factors (AF=4) may be preferred for studies including areas with elevated variance, but additional research is needed.

We examined leakage in single-subject analyses of the unprocessed HCP data, but the impacts in HCP preprocessed data were not examined. Analyzing fMRI data on the cortical surface with 2 mm surface smoothing and subcortical gray matter areas with parcel-constrained 2 mm volume smoothing in the manner advised by the WU-UMinn HCP consortium will decrease issues with false positives. Clearly, restricting an analysis to gray matter removes issues with aliasing artifacts in white matter. Secondly, performing 2 mm smoothing on the surface, which is less aggressive than volume smoothing, would reduce both smoothing leakage and slice leakage relative to our simulations, which use volume smoothing with FWHM equal to three times the voxel size. Todd et al. (2016) used 2 <u>mm</u> FWHM volume smoothing and found substantial slice leakage impacts with slice-GRAPPA reconstruction with multiband factors 4 and 6 combined with 2× in-plane acceleration. As discussed in Section 5, previous studies have found ICA can be used to identify artifacts related to the interaction between motion and SMS acquisition. Note that if an artifact associated with regional aliasing were identified with ICA, then the time course of the component would reflect the task activation. Then regressing out the time series of such a component would also remove the task variation from the region of true activation.

Another important question is how SMS acquisition impacts group-level analyses. In studies using the original slice-GRAPPA reconstruction, leakage could result in spurious clusters of activation. The impact may be reduced in a group analysis where subject variation in brain size and morphology results in subject-specific variation in the relative locations of aliased regions. However, when the activation region is relatively large, there may be a non-trivial overlap in the aliasing regions in MNI space and/or on the cortical surface, and then slice leakage may propagate to the group level. Additionally, group analyses often use larger amounts of smoothing (such as the 6 mm smoothing used in our single-subject analysis), which would increase the chance of aliasing artifacts propagating to the group level. In studies using HCP data, one can inspect group activation patterns in the LR versus RL encoded runs and see if differences are related to reversing the FOV shifts. It is also important to note that our approach to estimating slice leakage from the unprocessed HCP data is conservative in some respects. First, we do not count aliasing in regions that are not affected by FOV shifts, since these regions coincide in predicted RL and LR aliasing patterns and can not be distinguished from true positives. Secondly, the aliased regions corresponding to the FOV shifts often overlap, which further reduces the effective area over which our method can detect leakage. Thirdly, the online reconstruction algorithm includes corrections for spatial shifts due to B0 inhomogeneities, which may cause a discrepancy between our predicted aliasing patterns and the patterns of slice leakage in the “unprocessed” data. In simulations with similar effect sizes, there was evidence of leakage in only three-quarters of simulations, which was due to a combination of relatively little leakage at the small to moderate effect size and overlapping aliased regions in matched and mismatched PE directions.

There are limitations to our simulation study that should be noted. In real data applications, there are more structured differences between the single-band training data and the SMS data. Due to the short TR, T1 relaxation in SMS is incomplete, and the contrast between white and gray matter is greatly reduced (Glasser et al., 2013). This may affect kernel estimation and reconstruction accuracy. Our simulations capture impacts of AF on reconstruction accuracy due to noise amplification that arises from slice separation. To the extent that SMS data differ from the collapsed single-band data used in kernel estimation, we may overstate the sensitivity benefits of SMS. We have also not considered the impacts of subject motion on reconstruction. Slice-GRAPPA uses the same reconstruction kernel for all spatial frequencies in a given coil and given slice, see (S.6), and consequently head movement, particularly along the slice direction, can adversely impact image reconstruction accuracy. Reconstruction accuracy is also impacted by slice-specific Nyquist ghosting artifacts caused by spatially varying eddy currents (Barth et al., 2016), whereas our simulation approach effectively assumes there are no Nyquist ghosting artifacts. We also do not simulate higher-frequency artifacts, such that our SMS benefits are completely driven by the boost in effective sample size. Finally, we do not simulate serially correlated noise, such that the relative differences between AF = 8, AF = 4, and AF = 1 in sensitivity and specificity may be overstated.

## 7 Conclusion

Our contributions are the following:

- In simulations with a motor task, increasing the AF led to increased sensitivity, and slice-GRAPPA and split slice-GRAPPA had very similar sensitivity when combined with FOV/3 shifts.
- In slice-GRAPPA, 1 - specificity was inflated due to slice leakage and smoothing leakage. Slice leakage was greatly reduced by split slice-GRAPPA (leak block), although smoothing leakage resulted in inflated 1 - specificity. When using split slice-GRAPPA reconstruction, the slice leakage was adequately suppressed except at large effect sizes.
- Our findings suggest the amount of smoothing can be decreased in SMS acquisitions to increase spatial specificity where the boost in effective sample size helps offset the decrease in sensitivity.
- AF = 8 resulted in elevated measurement error in subcortical and some prefrontal regions and produced sharply defined regions of higher and lower measurement error in simulations. Similar patterns in residual variance were found in the unprocessed HCP motor task data and the ICA-FIX resting-state data, which also used AF = 8. In simulations, the issues were greatly reduced in AF = 4.
- We found evidence of slice leakage from a left-hand motor task contrast in smoothed unprocessed HCP data, which used slice-GRAPPA reconstruction. However, the detection of slice leakage was not ubiquitous across subjects.
- We did not analyze leakage in the ICA-FIX resting-state HCP data because preprocessing steps make it challenging to map aliased locations to the cortical surface and subcortical gray matter regions of grayordinate space. However, the less aggressive 2 mm surface smoothing for cortical data should greatly reduce the statistical impacts of leakage relative to the 6 mm volume smoothing used in our analysis of the unprocessed motor task data.

## 8 Acknowledgments

We thank Dr. Steen Moeller at the Center for Magnetic Resonance Research at the University of Minnesota for providing the single-band thirty-two channel reference data and for very helpful discussions. We also thank anonymous referees for their insights and suggestions, which have greatly improved the manuscript. Motor task and resting-state data were provided by the Human Connectome Project, WU-Minn Consortium (Principal Investigators: David Van Essen and Kamil Ugurbil; 1U54MH091657) funded by the 16 NIH Institutes and Centers that support the NIH Blueprint for Neuroscience Research; and by the McDonnell Center for Systems Neuroscience at Washington University. This material was based upon work partially supported by the NSF grant DMS-1127914 to the Statistical and Applied Mathematical Science Institute.

## Web Supplement to Impacts of Simultaneous Multislice Acquisition on Sensitivity and Specificity in fMRI

### A The Slice-GRAPPA algorithm

Here we give a formal description of slice-GRAPPA as created by Setsompop et al. (2012). Let *c* ∈ *1,…, C* index the coil at which we want to estimate the frequency-domain BOLD signal, where *C* = 32 in our application. Let ℓ ∈ {1,…, *Y*} and *k* ∈ {*1,…, X*} index the frequencies in the phase-encoding and read-out directions, respectively. Let *z* ∈ {*1,…, Z*} denote a slice in single-band space, where *z* = 72 in our application. Let t = 1,…, T denote the time point. Then let **S***^K^* ∈ ℂ*^C^*^×^*^Y^*^×^*^X^*^×^*^Z^*^×^*^T^* denote our full k-space data array, which we refer to as the spatiotemporal-by-coil k-space data, with entries 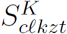. Our goal is to estimate **S***^K^* from multislice images. We denote the corresponding image-space representation as **S***^I^* ∈ ℂ*^C^*^×^*^Y^*^×^*^X^*^×^*^Z^*^×^*^T^*.

Let *A* denote the acceleration factor (number of simultaneously acquired slices). Let *M* = *Z*/*A* denote the number of multislice “packets.” Then let *m* ∈ {1,…, *M*} index a packet, and *m*(*z*) denote the packet that will be used to predict the BOLD signal at slice *z*:

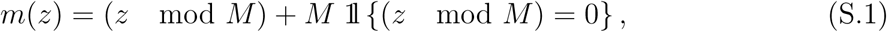
where (*z* mod *M*) denotes the remainder of *z*/*M*, defined as *z* for *z* < *M*. As an example, for *A* = 8 and *z* = 72, we have *M* = 9, and then slices 7, 16, 25, 34, 43, 52, 61, 70 are in packet 7.

Slice-GRAPPA uses data from all coils to improve prediction in a given coil. Let *h* = *1,…, C* denote the coils that will be used for reconstruction of the *c*th coil. Consider the spatial frequencies (*ℓ,k*) in the *m*th packet. Let 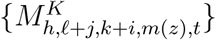 denote the aliased BOLD signal in the *m*th packet at the *t*th time point of the *h*th coil that will be used to predict the BOLD signal for the *c*th coil at location (*ℓ, k,z*) for the *t*th time point. Then let 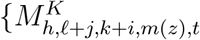, *h* ∈ (1,…, *C*}, *j* ∈ {−*J*,…, *J*}, *i* ∈ {*−I,…, I*}} denote the set of points that will be used to predict 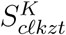. From a regression perspective, this set forms a vector of covariates that will be used to predict observation 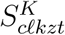.

Then define the coefficient 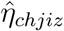, which characterizes the contribution of the *h*th coil at the location (*j*, *i*) away from an arbitrary focal location in the *z*th slice to the *c*th coil, which for now we take as given. Following Setsompop et al. (2012), the equation for predicting the signal 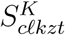 is

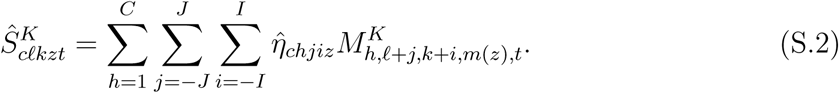

The set 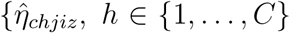, *j* = { − *J*,…, *J*}, *i* = {−*I*,…, *I*}} is called the “kernel” for the *c*th coil and *z*th slice because it specifies the weights for observations used to predict *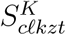*. Note the same kernel is used for predicting all *Y X* spatial frequencies for the *c*th coil and *z*th slice. Thus the set of kernels for reconstruction is a *C* × *C* × (2 *J* +1) × (2*I* + 1) × *Z* tensor. Typical values for the spatial frequency window size are *I* = 2 and *J* = 2 resulting in a 5-by-5 region (Uğurbil et al., 2013; Todd et al., 2016).

#### A.1 Estimating the kernel

The coefficients in (S.2) can be estimated from a single time point of single-band calibration data collected at the beginning of an fMRI session. Let *S_cℓkz0_* denote the single-band calibration data at the *z*th slice. Define 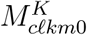 by adding the single-band data in the same manner as the multi-slice acquisition sequence:

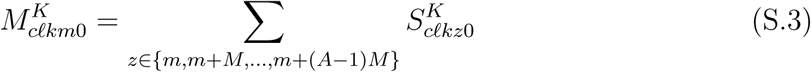
for each *m* ≥ *M*.

The coefficients are estimated using complex-valued least squares. Here we present a statistical model that can be used to derive the predictions in (S.2):

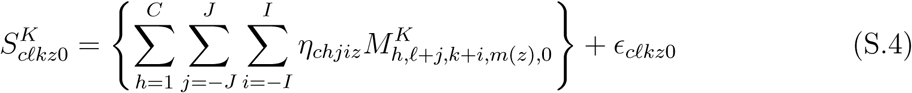
where 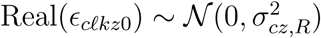 and 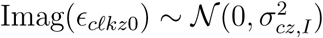.

For *J* < *ℓ* < *Y* − *J* and *I* < *k* < *X* − *I*, define the convolution matrix

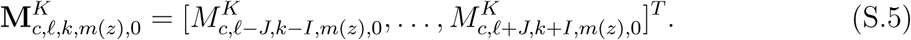

We can use all frequencies *ℓ* and *k* by allowing the indices to wrap such that for any *ℓ* ∈ {1*,…,Y* }*, j* ∈ {*−J, …, J*}*, k* ∈ {1*,…,X*}, and *i* ∈ {*−I,…,I*}, we revise (S.5) to have entries 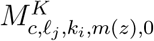 in which

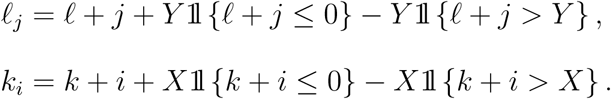

Next let 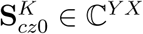 be the vectorized single-band data at slice *z*, where *YX* is the number of observations estimating the model parameters in (S.4). Then create

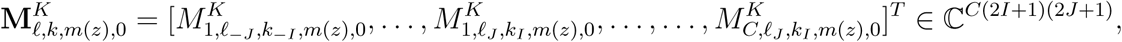
where ^“^*^T^*^”^ denotes the (ordinary) matrix transpose, and define

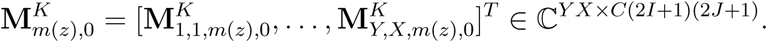

Define the vectorized kernel 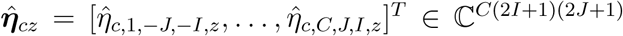. Then the coefficients are obtained using complex-valued least squares (e.g., Cauley et al. 2014):

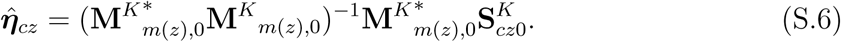
where ^“*”^ denotes the conjugate transpose. Note that the same design matrix is used for all coils and all slices in a multislice packet. After unaliasing the data in k-space, we apply the discrete inverse Fourier transform, and then take the sum-of-squares of the reconstructed coil data to generate the magnitude images.

### B Noise amplification in slice-GRAPPA

The standard deviation plots in the main manuscript are closely related to *g*-factors. In Figure S.1, we plot the ratio of the standard deviation from the accelerated data to the unaccelerated data, which is a measure of the *g*-factor when the signal intensities are equal (Robson et al., 2008).

**Figure S.1:**
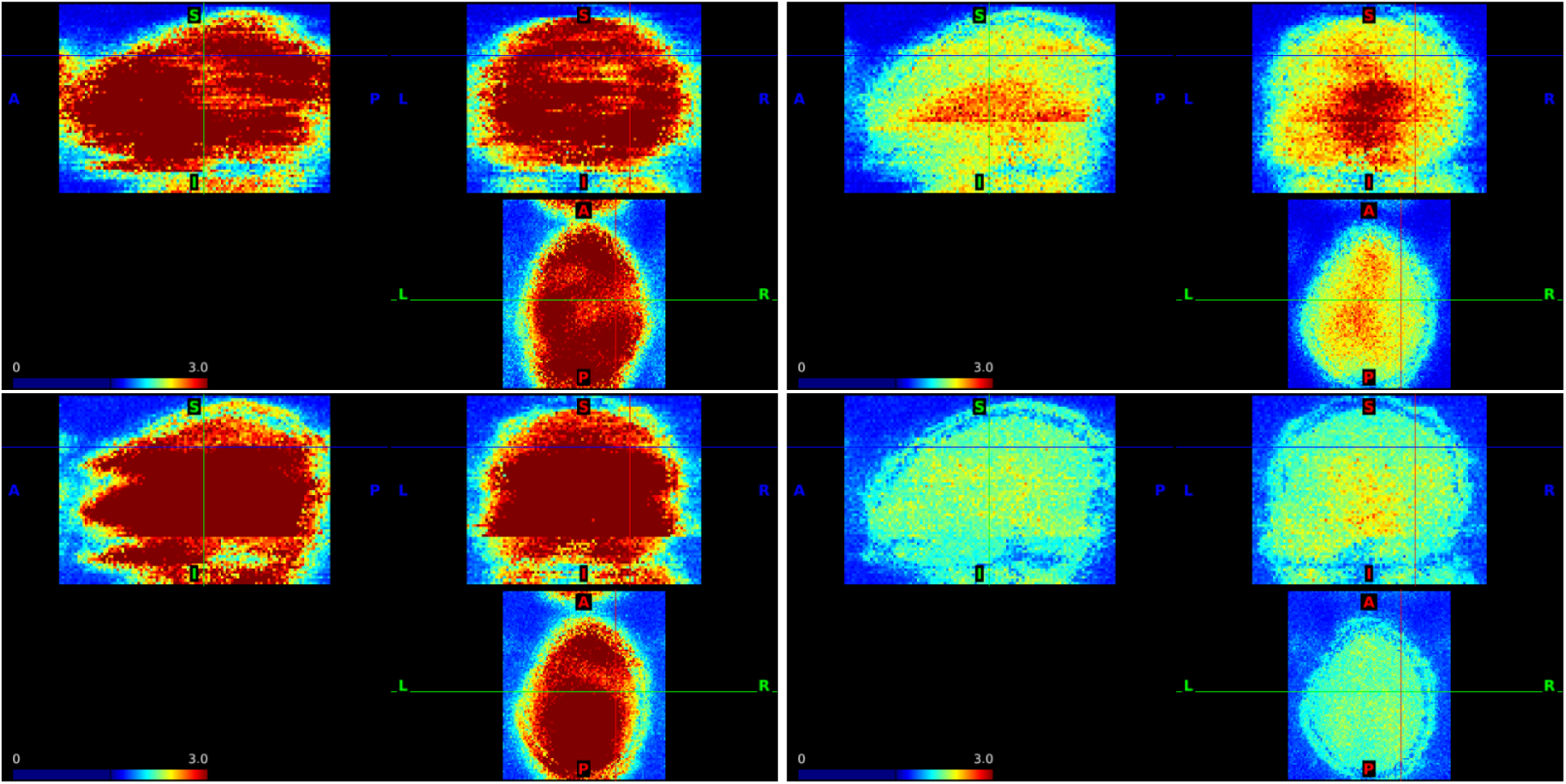
Estimates of *g*-factors using slice-GRAPPA based on the simulation design in Section 3.2 of the main manuscript. Top: AF = 8 with no shifts (left) and FOV/3 shifts (right). Bottom: AF = 4 with no shifts (left) and FOV/3 shifts (right).

### C Noise amplification in split slice-GRAPPA

The split slice-GRAPPA algorithm modifies the standard least squares problem by “splitting” the multiband data to increase the number of data points being predicted, such that the GRAPPA kernel is modified to fit zeros at the aliased locations; for details see Cauley et al. (2014).

**Figure S.2:**
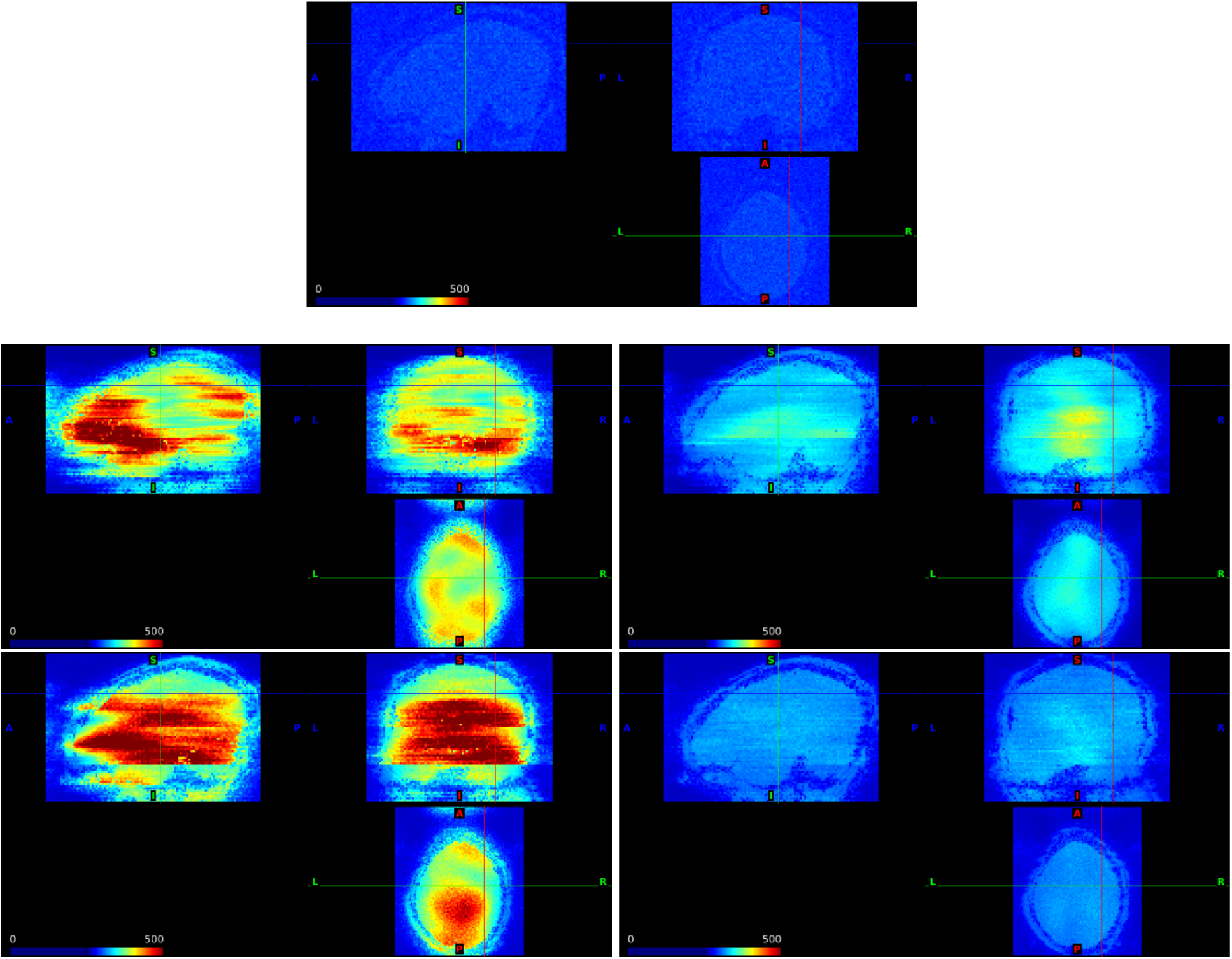
Noise amplification due to SMS using slice-GRAPPA reconstruction. Standard deviation of the residuals from the GLM fit to a simulation from Section 3.2 of the main manuscript with scaling factor = 1 and run duration = 480 s. Top: AF = 1; middle: AF = 8 with no FOV shifts (left) and FOV/3 shifts (right); bottom: AF = 4 with no FOV shifts (left) and FOV/3 shifts (right). This figure depicts the same information as Figure 2 of the main manuscript but uses the scale [0,500] instead of [0,250] to highlight differences between the no-CAIPI AF = 4 and AF = 8.

**Figure S.3:**
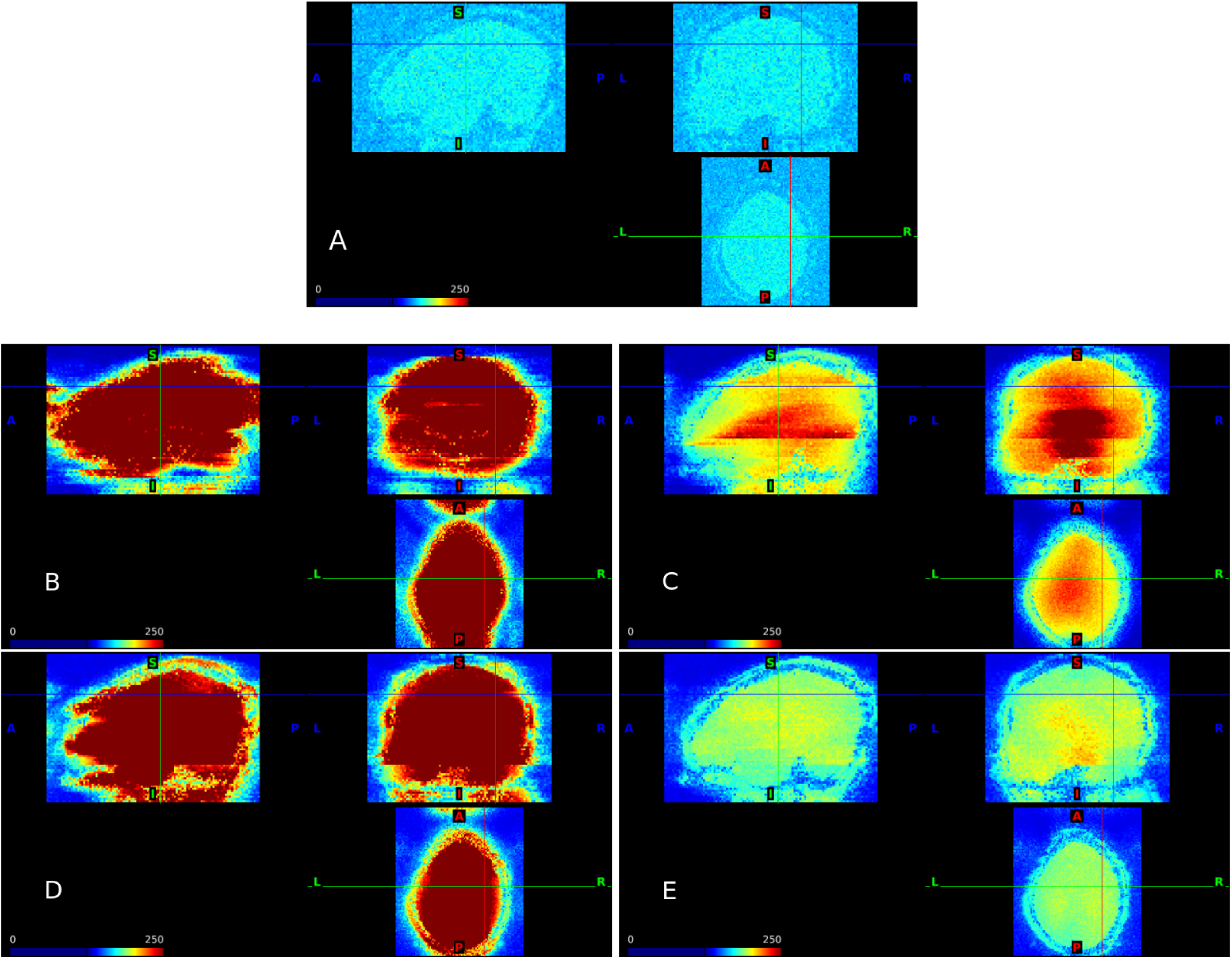
Noise amplification due to SMS using split slice-GRAPPA reconstruction (leak block). Standard deviation of the residuals from the GLM fit to a simulation from Section 3.2 of the main manuscript with scaling factor = 5 and run duration = 480 s. AF = 1 (A); AF = 8 with no FOV shifts (B) and FOV/3 shifts (C); AF = 4 with no FOV shifts (D) and FOV/3 shifts (E).

**Figure S.4:**
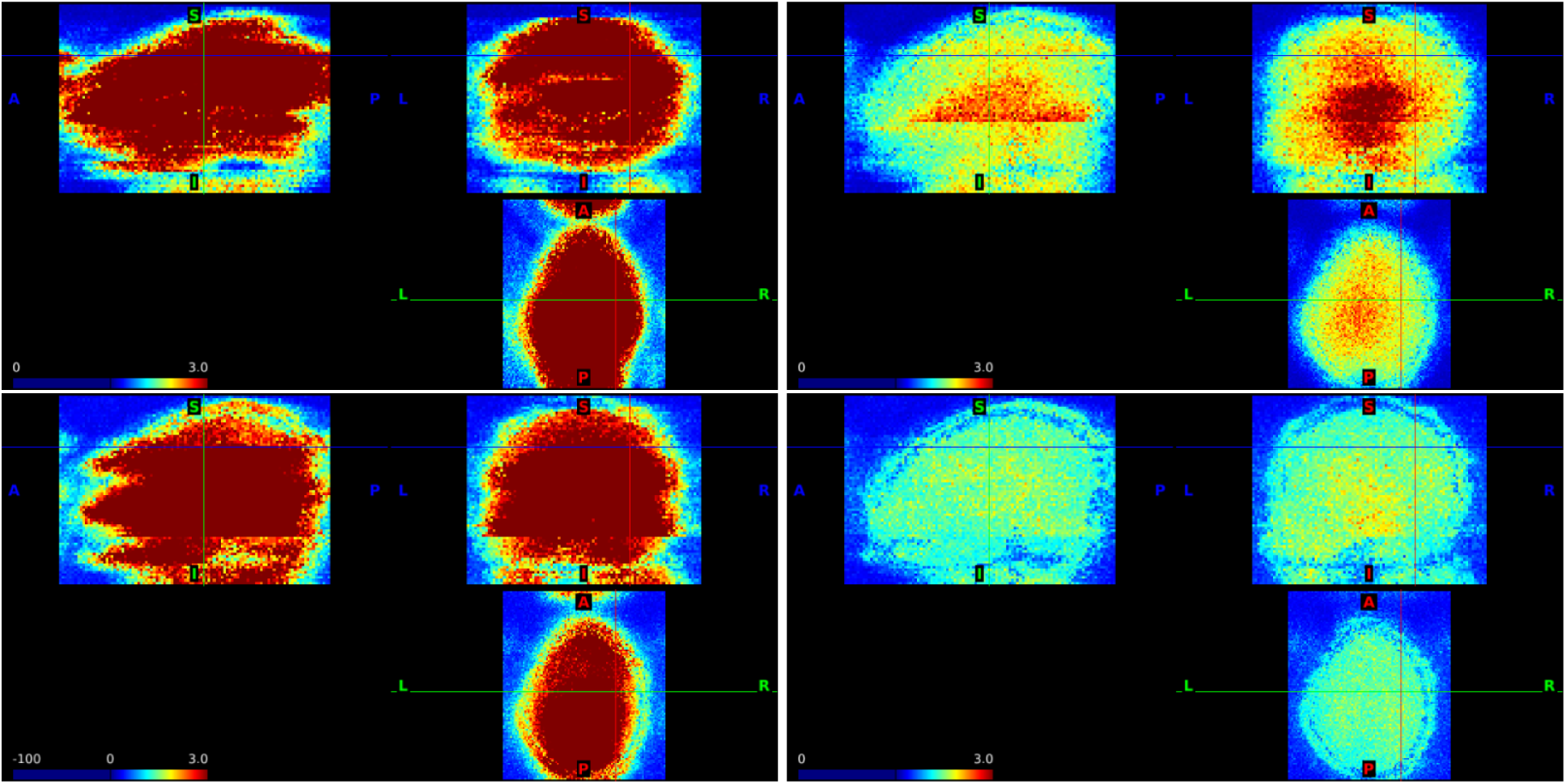
Estimates of *g*-factors using split slice-GRAPPA (leak block) based on the simulation design in Section 3.2 of the main manuscript. Top: AF = 8 with no shifts (left) and FOV/3 shifts (right). Bottom: AF = 4 with no shifts (left) and FOV/3 shifts (right).

### D Additional figure for sensitivity and specificity

**Figure S.5:**
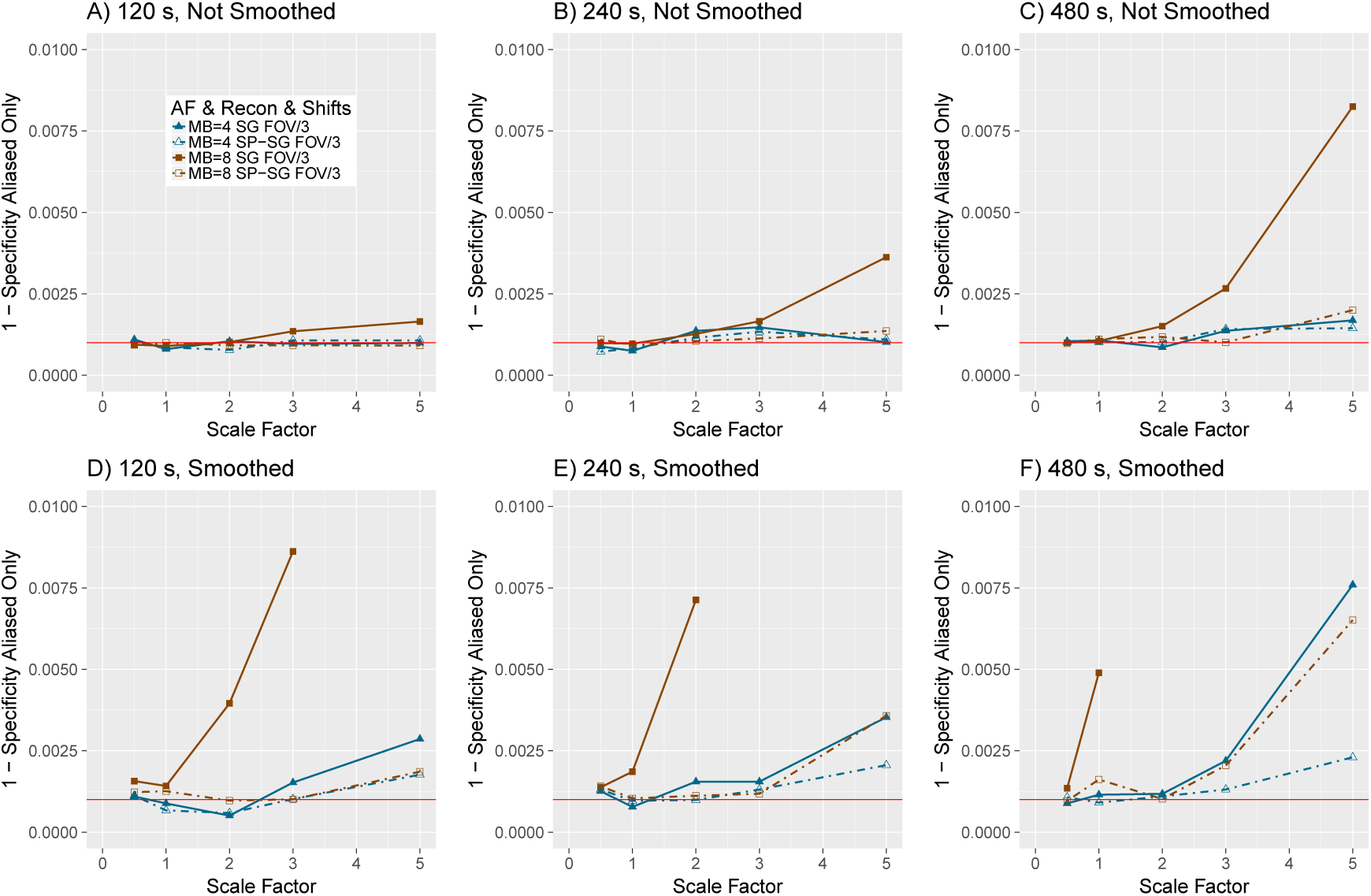
Zoomed in version of 1 - specificity (type 1 error rate) from Figures 3 and 4 of the main manuscript with no smoothing (A-C) and smoothing (D-F) for *α* = 0.001 and FOV/3 shifts. SG = slice-GRAPPA. SP-SG = split slice-GRAPPA. In panels D-F, values for AF = 8 and slice-GRAPPA reconstruction are not shown when they exceed 0.01. Scaling factors 0.5, 1, 2, 3, and 5 represent an average of 0.53%, 1.06%, 2.12%, 3.18%, and 5.30% change from baseline, respectively.

### E Synthetic multiband analysis

**Figure S.6:**
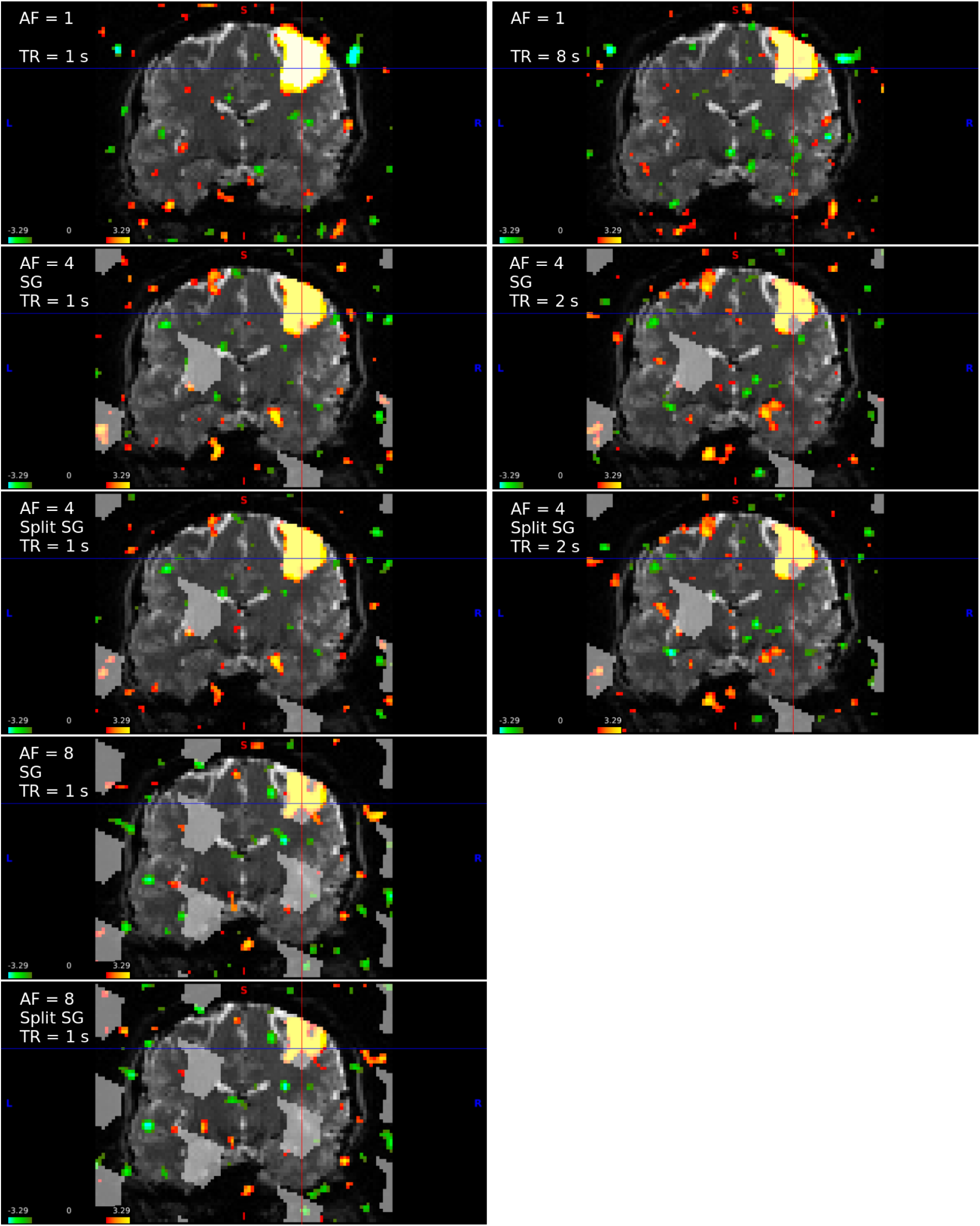
Synthetic simulation for scaling factor = 1. All figures are based on one singleband simulation, in which multiband data were synthetically generated. For TR = 1 s, differences between images are due to AF and reconstruction method. SG = slice-GRAPPA. Split SG = split slice-GRAPPA.

**Figure S.7:**
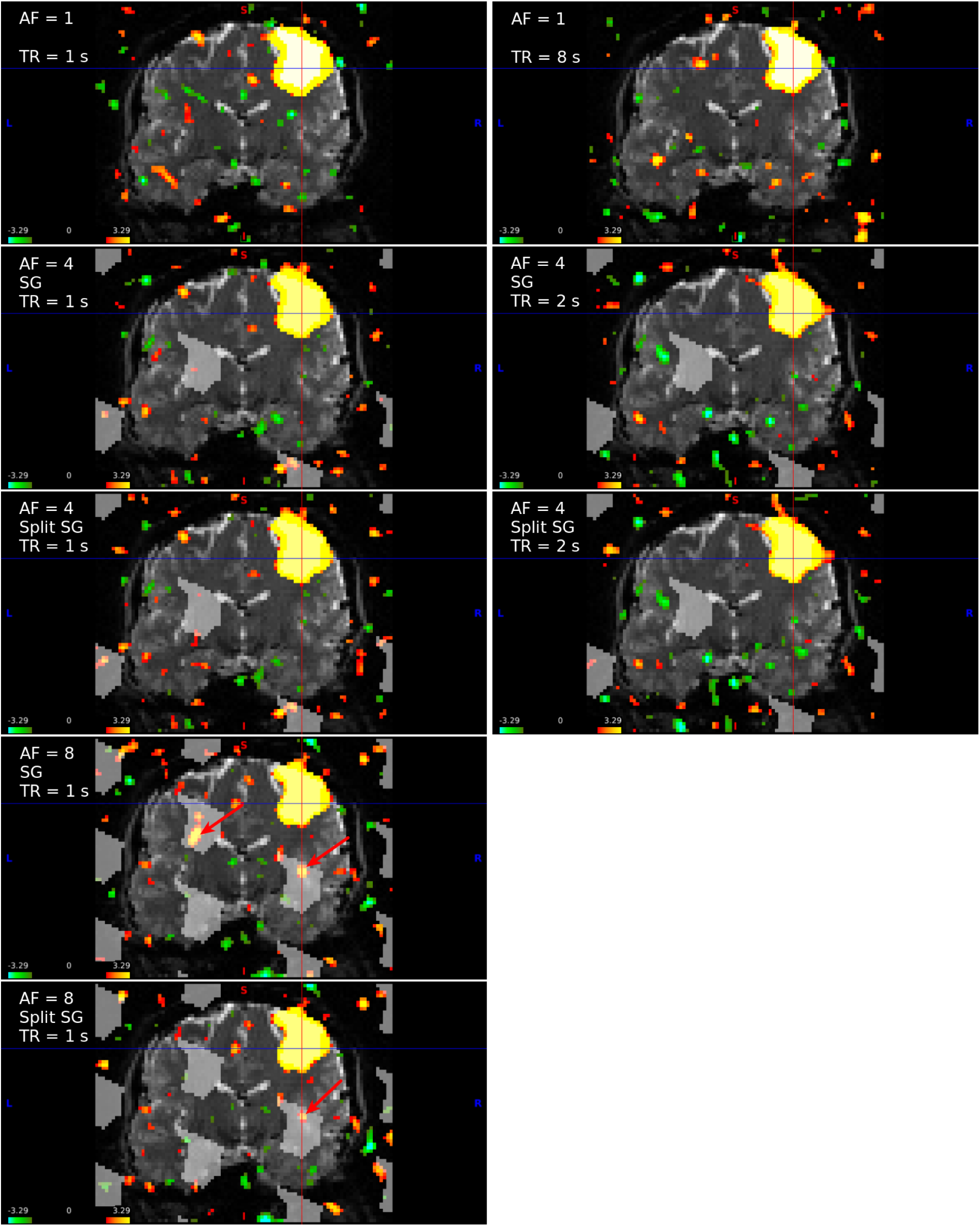
Synthetic simulation for scaling factor = 5. All figures are based on one singleband simulation, in which multiband data were synthetically generated. For TR = 1 s, differences between images are due to AF and reconstruction method. SG = slice-GRAPPA. Split SG = split slice-GRAPPA.

### F Additional information and results for the HCP analysis

**Table S.1:** Summary of acquisition parameters used by the HCP Consortium for the motor-task data. Additional details available in the HCP S1200 Release Appendix I (WU-Minn HCP Consortium, 2017).

| Parameters | Value |
| --- | --- |
| Sequence | Gradient-echo EPI |
| TR | 720 ms |
| TE | 33.1 ms |
| flip angle | 52 deg |
| Read direction | anterior-posterior |
| Phase-encoding direction | 1 run right-left, 1 run left-right |
| Slice acquisition | Inferior-superior |
| Position | L0.0 P3.0 H6.0 mm |
| Orientation | Transverse > Coronal - 20° |
| FOV | 208 mm × 180 mm × 144 mm |
| Voxel size | 2 × 2 × 2 mm<sup>3</sup> |
| Multiband factor | 8 |
| Blip-CAIPI Shift | FOV/3 in PE direction |
| In-plane acceleration | None |
| Echo spacing | 0.58 ms |
| BW | 2290 Hz/Px |
| Length per run | 3:41 |
| Frames per run | 284 |
| Volume data size unprocessed | 1.4 GB |
| Reconstruction | Slice-GRAPPA (no leak block) |

**Figure S.8:**
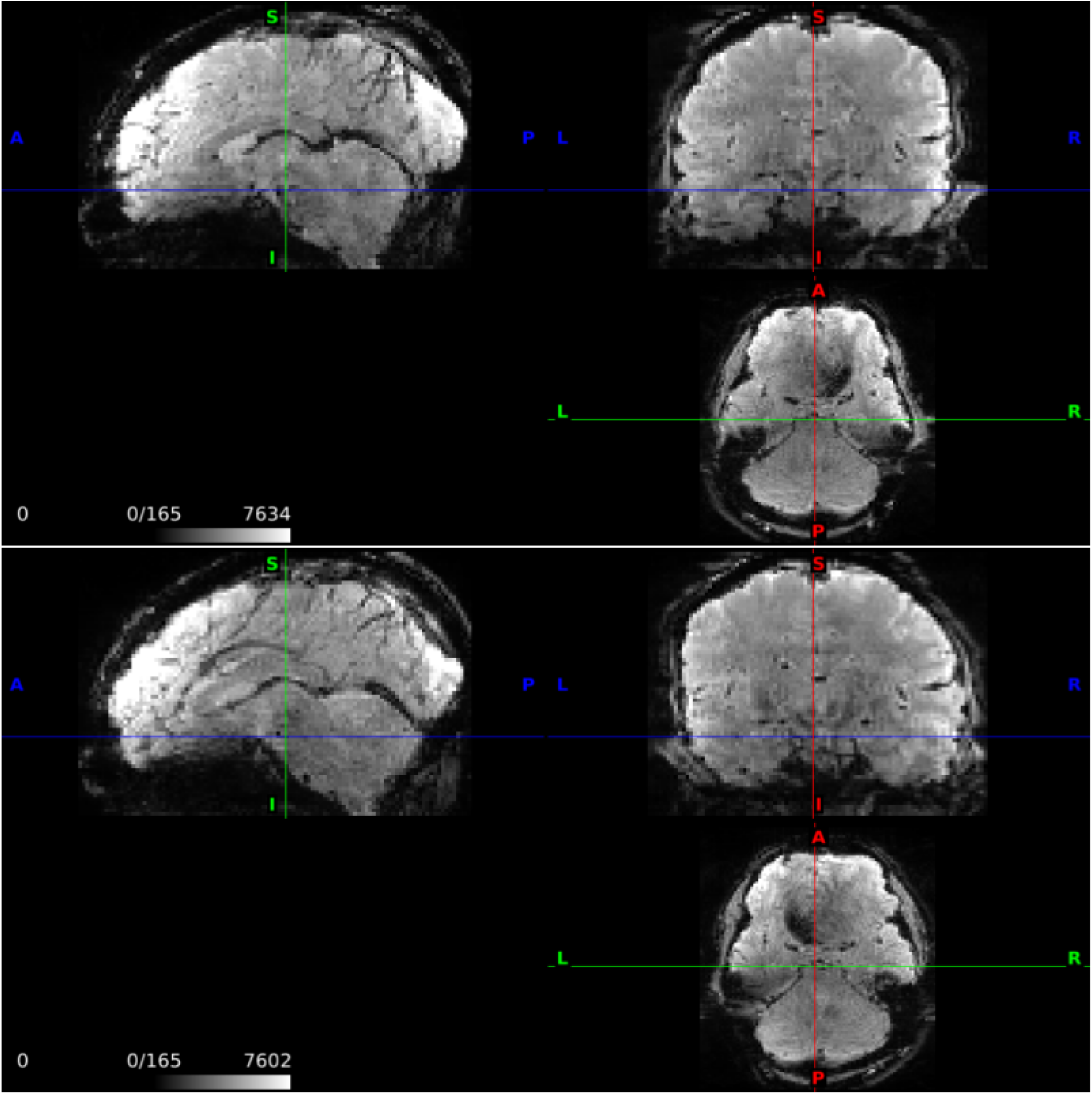
Mean image from the unprocessed HCP motor task fMRI data with RL encoding (top) and LR encoding (bottom) for subject 100206. The unprocessed HCP data generally have higher distortions from gradient non-linearities and EPI distortions as described in Glasser et al. (2013).

**Table S.2:** The proportion of voxels with *z* > 1.96 in the predicted aliased regions (matched) and in regions that are aliased using the opposite PE direction (mismatched). A larger proportion in *P_matched_* is considered evidence of slice leakage.

| | $P_{matched}$ | $P_{mismatched}$ | $Difference$ |
| --- | --- | --- | --- |
| 1 | 0.020 | 0.014 | 0.006 |
| 2 | 0.066 | 0.045 | 0.021 |
| 3 | 0.069 | 0.052 | 0.016 |
| 4 | 0.042 | 0.025 | 0.017 |
| 5 | 0.042 | 0.034 | 0.007 |
| 6 | 0.117 | 0.053 | 0.064 |
| 7 | 0.032 | 0.048 | -0.017 |
| 8 | 0.066 | 0.046 | 0.019 |
| 9 | 0.169 | 0.114 | 0.055 |
| 10 | 0.146 | 0.122 | 0.024 |
| 11 | 0.071 | 0.045 | 0.026 |
| 12 | 0.013 | 0.023 | -0.009 |
| 13 | 0.032 | 0.050 | -0.018 |
| 14 | 0.019 | 0.019 | -0.001 |
| 15 | 0.087 | 0.070 | 0.017 |
| 16 | 0.045 | 0.063 | -0.017 |
| 17 | 0.147 | 0.123 | 0.024 |
| 18 | 0.090 | 0.040 | 0.050 |
| 19 | 0.084 | 0.101 | -0.017 |
| 20 | 0.105 | 0.051 | 0.053 |
| 21 | 0.070 | 0.048 | 0.022 |
| 22 | 0.052 | 0.023 | 0.029 |
| 23 | 0.056 | 0.056 | 0.000 |
| 24 | 0.076 | 0.059 | 0.017 |
| 25 | 0.102 | 0.091 | 0.011 |
| 26 | 0.066 | 0.058 | 0.008 |
| 27 | 0.062 | 0.060 | 0.002 |
| 28 | 0.046 | 0.037 | 0.008 |
| 29 | 0.049 | 0.035 | 0.015 |
| 30 | 0.083 | 0.052 | 0.030 |
| 31 | 0.095 | 0.084 | 0.011 |
| 32 | 0.060 | 0.058 | 0.001 |
| 33 | 0.104 | 0.061 | 0.042 |
| 34 | 0.190 | 0.032 | 0.158 |
| 35 | 0.043 | 0.036 | 0.008 |
| 36 | 0.106 | 0.102 | 0.004 |
| 37 | 0.106 | 0.089 | 0.016 |
| 38 | 0.067 | 0.046 | 0.021 |
| 39 | 0.088 | 0.056 | 0.031 |
| 40 | 0.094 | 0.127 | -0.033 |
| 41 | 0.096 | 0.075 | 0.021 |
| 42 | 0.064 | 0.042 | 0.022 |
| 43 | 0.060 | 0.056 | 0.004 |
| 44 | 0.047 | 0.024 | 0.023 |
| 45 | 0.121 | 0.119 | 0.002 |
| 46 | 0.131 | 0.089 | 0.041 |
| 47 | 0.092 | 0.069 | 0.023 |
| 48 | 0.043 | 0.030 | 0.013 |
| 49 | 0.048 | 0.030 | 0.018 |
| 50 | 0.034 | 0.018 | 0.016 |
| 51 | 0.038 | 0.039 | -0.001 |
| 52 | 0.047 | 0.034 | 0.012 |
| 53 | 0.091 | 0.063 | 0.028 |
| 54 | 0.051 | 0.054 | -0.003 |
| 55 | 0.049 | 0.038 | 0.011 |
| 56 | 0.048 | 0.038 | 0.010 |
| 57 | 0.114 | 0.075 | 0.038 |
| 58 | 0.050 | 0.032 | 0.018 |
| 59 | 0.107 | 0.135 | -0.028 |
| 60 | 0.169 | 0.122 | 0.047 |
| 61 | 0.058 | 0.006 | 0.052 |
| 62 | 0.061 | 0.072 | -0.011 |
| 63 | 0.051 | 0.051 | -0.001 |
| 64 | 0.069 | 0.051 | 0.018 |
| 65 | 0.035 | 0.027 | 0.008 |
| 66 | 0.096 | 0.070 | 0.026 |
| 67 | 0.069 | 0.079 | -0.010 |
| 68 | 0.032 | 0.007 | 0.025 |
| 69 | 0.057 | 0.048 | 0.010 |
| 70 | 0.046 | 0.071 | -0.025 |
| 71 | 0.041 | 0.034 | 0.007 |
| 72 | 0.104 | 0.095 | 0.009 |
| 73 | 0.074 | 0.070 | 0.005 |
| 74 | 0.090 | 0.085 | 0.005 |
| 75 | 0.039 | 0.022 | 0.017 |
| 76 | 0.049 | 0.030 | 0.019 |
| 77 | 0.060 | 0.056 | 0.005 |
| 78 | 0.138 | 0.057 | 0.081 |
| 79 | 0.089 | 0.030 | 0.058 |
| 80 | 0.068 | 0.051 | 0.017 |
| 81 | 0.084 | 0.107 | -0.023 |
| 82 | 0.029 | 0.017 | 0.012 |
| 83 | 0.187 | 0.160 | 0.027 |
| 84 | 0.003 | 0.014 | -0.011 |
| 85 | 0.068 | 0.071 | -0.004 |
| 86 | 0.052 | 0.032 | 0.020 |
| 87 | 0.034 | 0.002 | 0.032 |
| 88 | 0.055 | 0.047 | 0.008 |
| 89 | 0.028 | 0.023 | 0.005 |
| 90 | 0.058 | 0.033 | 0.025 |
| 91 | 0.059 | 0.046 | 0.013 |
| 92 | 0.133 | 0.064 | 0.069 |
| 93 | 0.071 | 0.036 | 0.035 |
| 94 | 0.016 | 0.014 | 0.002 |
| 95 | 0.073 | 0.052 | 0.021 |
| 96 | 0.043 | 0.037 | 0.006 |
| 97 | 0.048 | 0.045 | 0.003 |
| 98 | 0.111 | 0.047 | 0.064 |

**Table S.3:**
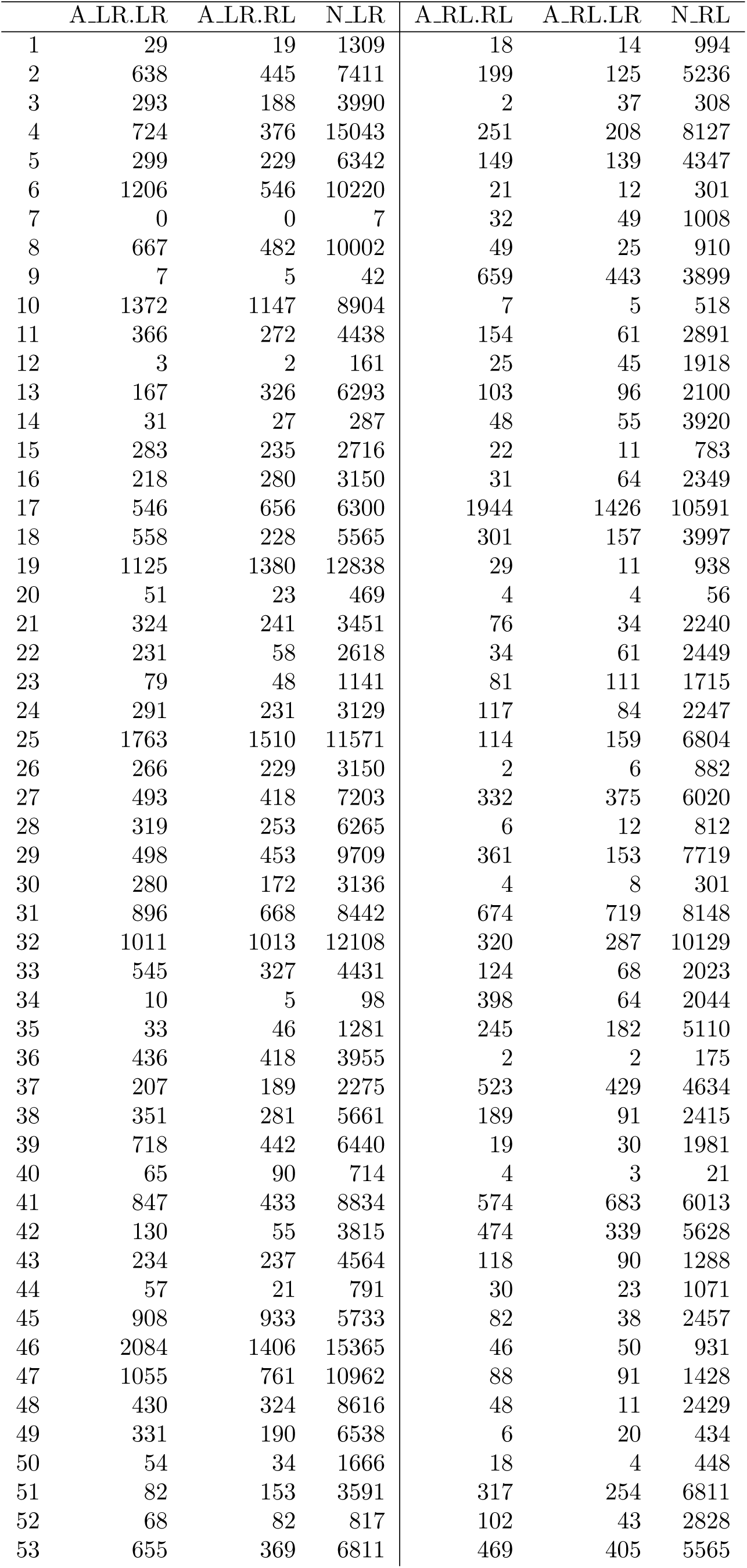
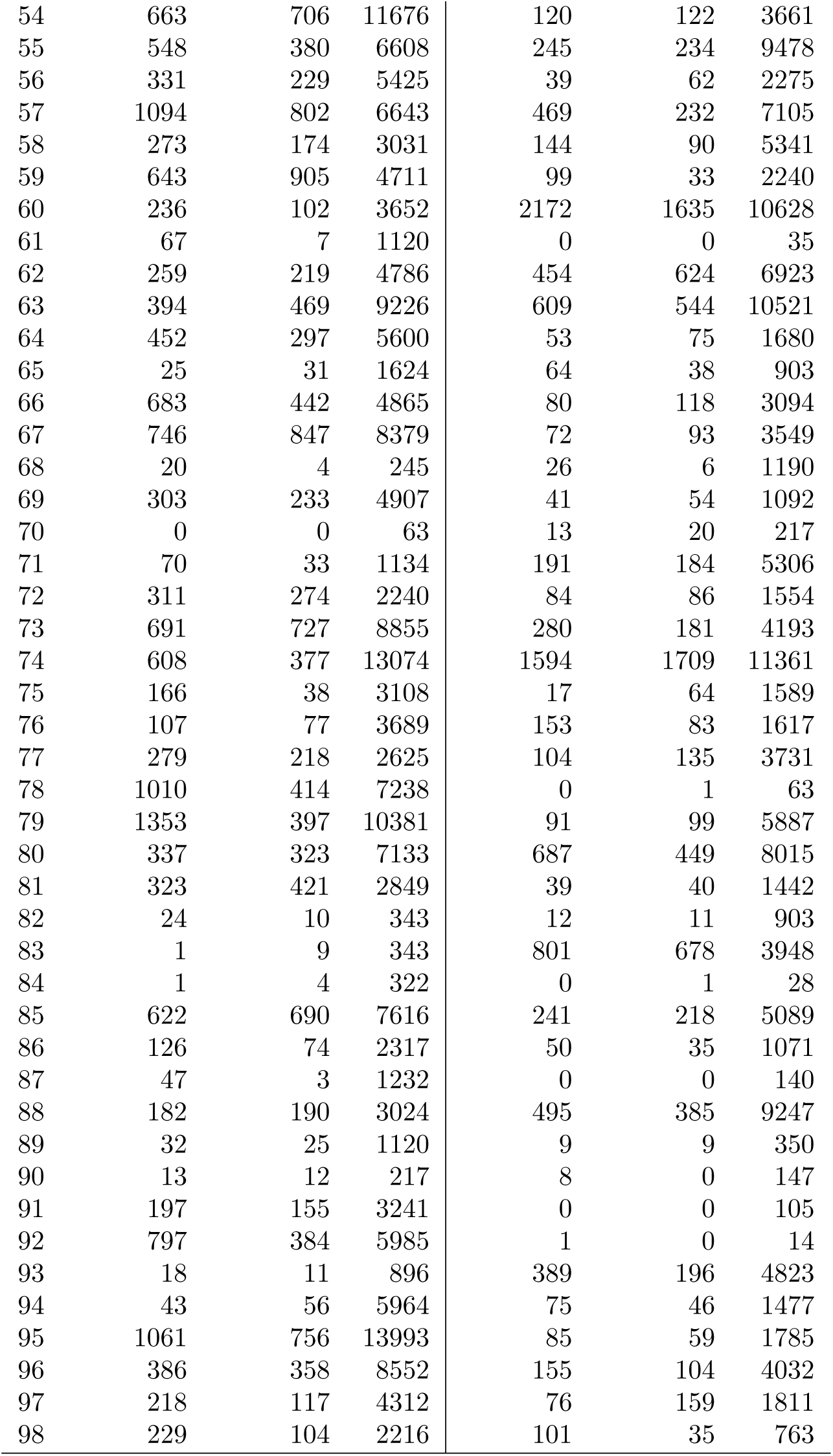
Counts of voxels with *z* > 1.96 used in the construction of the proportions in Table S.2.

## References

Aja-Fernández, S., Tristán-Vega, A., and Hoge, W. S. (2011). Statistical noise analysis in GRAPPA using a parametrized noncentral Chi approximation model. Magnetic resonance in medicine, 65(4):1195–1206.

Barth, M., Breuer, F., Koopmans, P. J., Norris, D. G., and Poser, B. A. (2016). Simultaneous multislice (SMS) imaging techniques. Magnetic resonance in medicine, 75(1):63–81.

Bilgic, B., Gagoski, B. A., Cauley, S. F., Fan, A. P., Polimeni, J. R., Grant, P. E., Wald, L. L., and Setsompop, K. (2015). Wave-CAIPI for highly accelerated 3D imaging. Magnetic resonance in medicine, 73(6):2152–2162.

Boyacioğlu, R., Schulz, J., Koopmans, P. J., Barth, M., and Norris, D. G. (2015). Improved sensitivity and specificity for resting state and task fMRI with multiband multi-echo EPI compared to multi-echo EPI at 7T. Neuroimage, 119:352–361.

Breteler, M. M., Stöcker, T., Pracht, E., Brenner, D., and Stirnberg, R. (2014). MRI in the Rhineland study: A novel protocol for population neuroimaging. Alzheimer’s & Dementia: The Journal of the Alzheimer’s Association, 10(4):P92.

Breuer, F. A., Blaimer, M., Heidemann, R. M., Mueller, M. F., Griswold, M. A., and Jakob, P. M. (2005). Controlled aliasing in parallel imaging results in higher acceleration (CAIPIR-INHA) for multi-slice imaging. Magnetic resonance in medicine, 53(3):684–691.

Bruce, I. P. and Rowe, D. B. (2014). Quantifying the statistical impact of GRAPPA in fcMRI data with a real-valued isomorphism. IEEE transactions on medical imaging, 33(2):495–503.

Cauley, S. F., Polimeni, J. R., Bhat, H., Wald, L. L., and Setsompop, K. (2014). Inter-slice leakage artifact reduction technique for simultaneous multislice acquisitions. Magnetic Resonance in Medicine, 72(1):93–102.

Chen, J. E. and Glover, G. H. (2017). Modeling serial correlations of fmri time series collected by faster TRs. 25th Annual Meeting of ISMRM.

Chen, L., Vu, A. T., Xu, J., Moeller, S., Ugurbil, K., Yacoub, E., and Feinberg, D. A. (2015). Evaluation of highly accelerated simultaneous multi-slice EPI for fMRI. Neuroimage, 104:452–459.

Demetriou, L., Kowalczyk, O. S., Tyson, G., Bello, T., Newbould, R. D., and Wall, M. B. (2016). A comprehensive evaluation of multiband-accelerated sequences and their effects on statistical outcome measures in fMRI. bioRxiv, page 076307.

Feinberg, D. A., Moeller, S., Smith, S. M., Auerbach, E., Ramanna, S., Glasser, M. F., Miller, K. L., Ugurbil, K., and Yacoub, E. (2010). Multiplexed echo planar imaging for sub-second whole brain fMRI and fast diffusion imaging. PloS one, 5(12):e15710.

Feinberg, D. A. and Setsompop, K. (2013). Ultra-fast MRI of the human brain with simulta-neous multi-slice imaging. Journal of magnetic resonance, 229:90–100.

Glasser, M. F., Smith, S. M., Marcus, D. S., Andersson, J. L., Auerbach, E. J., Behrens, T. E., Coalson, T. S., Harms, M. P., Jenkinson, M., Moeller, S., et al. (2016). The human connectome project’s neuroimaging approach. Nature Neuroscience, 19(9):1175–1187.

Glasser, M. F., Sotiropoulos, S. N., Wilson, J. A., Coalson, T. S., Fischl, B., Andersson, J. L., Xu, J., Jbabdi, S., Webster, M., Polimeni, J. R., et al. (2013). The minimal preprocessing pipelines for the Human Connectome Project. NeuroImage, 80:105–124.

Griffanti, L., Douaud, G., Bijsterbosh, J., Evangelisti, S., Alfaro-Almagro, F., Glasser, M. F., Duff, E. P., Fitzgibbon, S., Westphal, R., Carone, D., et al. (2016). Hand classification of fMRI ICA noise components. NeuroImage.

Griffanti, L., Salimi-Khorshidi, G., Beckmann, C. F., Auerbach, E. J., Douaud, G., Sexton, C. E., Zsoldos, E., Ebmeier, K. P., Filippini, N., Mackay, C. E., et al. (2014). ICA-based artefact removal and accelerated fMRI acquisition for improved resting state network imaging. NeuroImage, 95:232–247.

Miller, K. L., Alfaro-Almagro, F., Bangerter, N. K., Thomas, D. L., Yacoub, E., Xu, J., Bartsch, A. J., Jbabdi, S., Sotiropoulos, S. N., Andersson, J. L., et al. (2016). Multimodal population brain imaging in the UK Biobank prospective epidemiological study. Nature Neuroscience.

Moeller, S., Xu, J., Auerbach, E., Yacoub, E., and Ugurbil, K. (2012). Signal leakage (L-factor) as a measure for parallel imaging performance among simultaneously multi-slice (SMS) excited and acquired signals. In Proceedings of the 20th Annual Meeting of ISMRM, Melbourne, Australia, page 519.

Moeller, S., Yacoub, E., Olman, C. A., Auerbach, E., Strupp, J., Harel, N., and Uğurbil, K. (2010). Multiband multislice GE-EPI at 7 tesla, with 16-fold acceleration using partial parallel imaging with application to high spatial and temporal whole-brain fMRI. Magnetic Resonance in Medicine, 63(5):1144–1153.

Nunes, R., Hajnal, J., Golay, X., and Larkman, D. (2006). Simultaneous slice excitation and reconstruction for single shot EPI. In Proceedings of the 14th annual meeting of ISMRM, Seattle, Washington, USA, page 293.

Penny, W. D., Friston, K. J., Ashburner, J. T., Kiebel, S. J., and Nichols, T. E. (2007). Statistical parametric mapping: the analysis of functional brain images. Academic press.

Pfeuffer, J., de Moortele, V., Ugurbil, K., Hu, X., Glover, G. H., et al. (2002). Correction of physiologically induced global off-resonance effects in dynamic echo-planar and spiral functional imaging. Magnetic Resonance in Medicine, 47(2):344–353.

Poldrack, R. A., Mumford, J. A., and Nichols, T. E. (2011). Handbook of functional MRI data analysis. Cambridge University Press.

Poser, B. A. and Setsompop, K. (2017). Pulse sequences and parallel imaging for high spatiotemporal resolution MRI at ultra-high field. NeuroImage.

Preibisch, C., Bührer, M., Riedl, V., et al. (2015). Evaluation of multiband EPI acquisitions for resting state fMRI. PloS one, 10(9):e0136961.

Pruessmann, K. P., Weiger, M., Scheidegger, M. B., Boesiger, P., et al. (1999). SENSE: sensitivity encoding for fast MRI. Magnetic resonance in medicine, 42(5):952–962.

Risk, B. B., Matteson, D. S., Spreng, R. N., and Ruppert, D. (2016). Spatiotemporal mixed modeling of multi-subject task fMRI via method of moments. NeuroImage.

Robson, P. M., Grant, A. K., Madhuranthakam, A. J., Lattanzi, R., Sodickson, D. K., and McKenzie, C. A. (2008). Comprehensive quantification of signal-to-noise ratio and g-factor for image-based and k-space-based parallel imaging reconstructions. Magnetic resonance in medicine, 60(4):895–907.

Salimi-Khorshidi, G., Douaud, G., Beckmann, C. F., Glasser, M. F., Griffanti, L., and Smith, S. M. (2014). Automatic denoising of functional MRI data: combining independent component analysis and hierarchical fusion of classifiers. Neuroimage, 90:449–468.

Setsompop, K., Gagoski, B. A., Polimeni, J. R., Witzel, T., Wedeen, V. J., and Wald, L. L. (2012). Blipped-controlled aliasing in parallel imaging for simultaneous multislice echo planar imaging with reduced g-factor penalty. Magnetic Resonance in Medicine, 67(5):1210–1224.

Smith, S. M., Beckmann, C. F., Andersson, J., Auerbach, E. J., Bijsterbosch, J., Douaud, G., Duff, E., Feinberg, D. A., Griffanti, L., Harms, M. P., et al. (2013). Resting-state fMRI in the human connectome project. Neuroimage, 80:144–168.

Todd, N., Josephs, O., Zeidman, P., Flandin, G., Moeller, S., and Weiskopf, N. (2017). Functional sensitivity of 2D simultaneous multi-slice echo-planar imaging: effects of acceleration on g-factor and physiological noise. Frontiers in Neuroscience, 11.

Todd, N., Moeller, S., Auerbach, E. J., Yacoub, E., Flandin, G., and Weiskopf, N. (2016). Evaluation of 2D multiband EPI imaging for high-resolution, whole-brain, task-based fMRI studies at 3T: Sensitivity and slice leakage artifacts. NeuroImage, 124:32–42.

Uğurbil, K., Xu, J., Auerbach, E. J., Moeller, S., Vu, A. T., Duarte-Carvajalino, J. M., Lenglet, C., Wu, X., Schmitter, S., Van de Moortele, P. F., et al. (2013). Pushing spatial and temporal resolution for functional and diffusion MRI in the human connectome project. NeuroImage, 80:80–104.

Worsley, K. J., Liao, C., Aston, J., Petre, V., Duncan, G., Morales, F., and Evans, A. (2002). A general statistical analysis for fMRI data. NeuroImage, 15(1):1–15.

WU-Minn HCP Consortium (2015). 900 subjects data release: Reference manual.

Xu, J., Moeller, S., Auerbach, E. J., Strupp, J., Smith, S. M., Feinberg, D. A., Yacoub, E., and Uğurbil, K. (2013). Evaluation of slice accelerations using multiband echo planar imaging at 3T. NeuroImage, 83:991–1001.

Yarkoni, T., Poldrack, R. A., Nichols, T. E., Van Essen, D. C., and Wager, T. D. (2011). Large-scale automated synthesis of human functional neuroimaging data. Nature methods, 8(8):665–670.

Zhu, K., Dougherty, R., Pauly, J., and Kerr, A. (2014). Multislice acquisition with incoherent aliasing (mica). In Proceedings of the 22nd Annual Meeting of ISMRM, Milan, Italy.

## References

Cauley, S. F., Polimeni, J. R., Bhat, H., Wald, L. L., and Setsompop, K. (2014). Interslice leakage artifact reduction technique for simultaneous multislice acquisitions. Magnetic Resonance in Medicine, 72(1):93–102.

WU-Minn HCP Consortium (2017). 1200 subjects data release reference manual.

